# *Predominance of clonal propagation* conceals extinction risks of the highly endangered floodplain herb *Cnidium dubium*

**DOI:** 10.1101/2021.10.22.465465

**Authors:** Ilona Leyer, Birgit Ziegenhagen, Christina Mengel, Eva Mosner, Sascha Liepelt

## Abstract

Habitat loss and degradation due to human-induced landscape alterations are considered to be a major threat to biodiversity. The decline of biodiversity may occur with a time delay leading to a so called extinction debt. Therefore, determining extinction risks and conservation status is not always straightforward. Several life history traits might play a role for the accumulation of an extinction debt. Thus, perennial plant species capable of vegetative propagation might be able to persist temporarily in degraded habitats even though sexual and evolutionary processes are effectively halted.

We studied *Cnidium dubium*, which occurs in scattered patches along river corridors in Central Europe and is critically endangered in Germany. It is a perennial species which is able to propagate clonally. Our aims were to reconstruct demographic processes regarding clonal propagation and gene flow along 400 km of river stretch and with respect to the position in the flooplain, i.e. before or behind dykes. We also wanted to determine whether there is evidence for an extinction debt in *C. dubium* and to use our insights for conservation recommendations.

For this, we used nuclear microsatellites and maternally inherited chloroplast DNA markers and applied a systematic grid based sampling strategy for small scale geographic structures.

We observed a high level of clonal propagation. In 935 analysed plants we observed only 121 different genotypes and of 50 studied patches of *C. dubium* the majority (31 patches) consisted of one single genotype each. Patch size and position were correlated with clonal diversity. Large patches and patches behind dykes exhibited higher clonal diversity. There was no evidence for a large scale genetic substructuring of the study area and no differences in overall genetic diversity between upstream and downstream patches as well as between patches before and behind the dykes. High levels of heterozygosity and a high number of 18 chloroplast DNA haplotypes togetherwith a slightly elevated inbreeding coefficient (Fis) point toward a high level of ancestral polymorphism in an out of equilibrium population due to high levels of clonal propagation and low levels of gene flow and recombination. Therefore, we assume that an extinction debt is present in C. dubium. As a management strategy, we propose to transplant ramets between multiple patches to increase the number of mating partners and therefore restore sexual reproduction.

## Introduction

Human-induced alterations in landscape structure and concomitant loss of natural and semi-natural habitats pose a major threat to biodiversity (Tilman et al. 2001, Tscharntke et al. 2011). Habitat loss can involve a concurrent decline in habitat area and connectivity as well as in the quality of habitat patches (Fahrig 2003, Sang et al. 2010). Reduced habitat area and isolation is accompanied by decreased population sizes and restricted gene flow within species (Honnay et al. 2005, Duwe et al. 2017, Lee et al. 2018). Together with deteriorating habitat quality, survival and reproduction of individuals are threatened and their fitness is reduced (Colling & Matthies 2006, Mortelliti et al. 2010). Decline of biodiversity in response to habitat loss can be an immediate evident, but extinction events might also occur with a time delay called “relaxation time” (Diamond et al. 1972) due to the phenomenon of extinction debt (Tilman et al. 1994, Hanski & Ovaskainen 2002). Extinction debt means that species in a local community are doomed to extinction due to altered environmental, e.g. habitat conditions, but the actual extinction event has not yet occurred since species can persist for a time in small, isolated and degraded habitats (Kuussaari et al. 2009, Krauss et al. 2010). Extinction debt can lead to an underestimation of actual threats to biodiversity (Tilman et al. 1994, Bulman et al. 2007) as often only a simple count of species is used to evaluate conservation needs of habitats (Helm et al. 2005). Likewise, the vulnerability of individual species can be underestimated due to an extinction debt if the population size rather reflects the former area, connectivity and quality of the habitat rather than the current one (Tepedino 2012, Hylander & Ehrlén 2013). Thus, this phenomenon can easily remain unrecognized and should be taken into account when planning restoration schemes.

Although information is scarce on the influence of plant species traits on extinction debt, empirical evidence suggests that time-delayed extinctions are more likely to occur in long-lived species compared to short-lived ones. Thus, it can be assumed that perennial rather than annual plants as well as tree species may carry an extinction debt (Kuussaari et al. 2009). Furthermore, there is evidence that isolated populations of clonally propagating species can persist long after a habitat fragmentation event (Honnay et al. 2005). Reasons could be that they are more buffered against the heterogeneity of their habitats due to the reallocation of resources and division of labour among ramets of a genet. The probability of genet death can be reduced by spreading the risk over multiple ramets (Stuefer et al. 1996, Pennings & Callaway 2000, Honnay & Bossuyt 2005). Furthermore, there is evidence that vegetative reproduction can be maintained in habitats in which sexual reproduction is prevented, e.g. where ecological conditions become unfavorable for seed set, seed germination, or seedling establishment (Lindborg & Eriksson 2004). Thus, resulting population structure could be rather shaped by the former landscape than by the degree of fragmentation (Young et al. 1996, Honnay et al. 2005, Llorens et al. 2018). Remnant populations of long-lived clonal plants might therefore appear large and viable, but might be doomed to extinction in the long run. Despite the benefits of clonal propagation, prolonged clonal growth can also be negative as it can limit the outcome of sexual reproduction due to e.g. limited resource allocation to flowering and seed production and the interference of vegetative reproduction with pollination and mating (Vallejo-Marín et al. 2010, Barrett 2015). Furthermore, the size and longevity of clonal populations could be associated with the accumulation of somatic mutations due to high numbers of somatic cell divisions in old clones potentially leading to degeneration (Bobiwash et al. 2013, Barrett 2015). These aspects illustrate clearly that knowledge of clonal diversity and the extent of clonal structures is a necessary prerequisite to assess the threat to clonal plant populations and their chances of survival.

Floodplain habitats with their peculiar species communities belong to the most altered and fragmented ecosystems around the world. Many plant species and their habitats confined to river corridors have strongly declined in the last centuries due to hydrological alterations through river regulation by dams for e.g. navigation and hydroelectric power production (Lehner et al. 2011) as well as floodplain fragmentation by dykes (also called levees) for flood protection (Leyer 2005). Dykes have led to a dramatic decrease in the actively flooded area of nearly all river systems in Central Europe. In Germany, all major rivers (e.g. Rhine, Elbe, Oder, Danube) have lost more than two thirds of their active floodplains (BMU and BFN 2009). Behind the dykes, in the inactive floodplain, flooding and flow induced disturbances are prevented leading to accelerated settlement activities and land use intensification. Due to the poor state of these habitats and many of its representative plant species, they are part of strong conservation and restoration efforts (Mosner et al. 2012, Schindler et al. 2016).

As explained above, the assessment of threats to clonal species by floodplain fragmentation and deterioration is a challenge. However, as for population genetic effects, not only clonal diversity and the extent of clonal structures but also genetic diversity and differentiation affected by gene flow have to be considered. It is well known, that unidirectional water flow can link plant populations over long distances due to water dispersal (hydrochory) (Kudoh & Whigham 1997, 2001; Kondo et al. 2009). This can lead to low genetic divergence along the river (Jacquemyn et al. 2006, Hu et al. 2010). Since seed dispersal by water is unidirectional, in some studies an increase of genetic diversity downriver could be observed (Nilsson et al. 2010, Schleuning et al. 2011). These processes can only come into action in the active floodplain, not in the floodplain behind the dykes, where water flow is prevented. For sites in the inactive floodplain profound effects on the plant population level can be expected. However, knowledge regarding this topic is rare (Nilsson et al. 2010, but see Mosner et al. 2012 for *Salix viminalis*).

In this study, we applied microsatellite and cpDNA markers to infer clonal patterns as well as gene flow and water dispersal processes in *Cnidium dubium* (Schkuhr) Thell. along a 400 km course of the Elbe River, Germany. *C. dubium* is a hemicryptophytic species of the Apiaceae. In Central Europe, it occurs predominantly in floodplain meadows along the corridors of large rivers (Vent & Benkert 1984, Burkart 2001). Due to river regulation and floodplain fragmentation by dykes with subsequent intensification of land use as well as abandonment and drainage of floodplain and wetland meadows, populations are strongly declining (BfN 2017). Interestingly and as an advantageous setting for our study the species occurs equally distributed in both the active and inactive floodplain of the Elbe River. *C. dubium* is listed as critically endangered in the Red List of vascular plants in Germany (RL status 2). Its habitats are listed in Annex I of the EU-Habitat Directive (code 6440: Alluvial meadows of river valleys of the alliance *Cnidion dubii*). In Germany, they are acutely threatened with extinction (RL 1, Finck et al. 2017). The largest remnants of *Cnidium* meadows in central Europe are to be found in the floodplains of the river Elbe, but their area has decreased continuously in recent decades as a result of changing land-use practices and trophic conditions.

We used nuclear microsatellite markers in order to unravel small-scale patterns of clonal structures as well as genetic diversity and differentiation along the river taking floodplain type (active/inactive) as well as the size of *Cnidium* patches (small/large) into account. Moreover, we applied chloroplast DNA markers to provide important information about the demographic history of the studied populations because their distribution is linked to seed dispersal events. Including both chloroplast haplotypic and nuclear microsatellite information we inferred historic and recent gene flow processes and linked them to discovered clonal structures.

Specifically, we aimed to answer the following questions:

1. How are clonal and genetic structures spatially organized within *C. dubium* patches and do the observed patterns point towards an extinction debt?
2. Are there differences in genetic diversity and differentiation regarding active and inactive floodplain as well as along the course of the Elbe River and do results suggest that water dispersal and other gene flow processes shape large scale population genetic structure?
3. Which conclusions can be drawn and what are promising measures for successful conservation and restoration of *C. dubium* populations and other endangered clonal plant species in floodplain ecosystems?

## Methods

### Sampling

Leaf material was sampled from *C. dubium* patches in May and October 2012. A sample is congruent with a leaf of a shoot and hereinafter termed “ramet” to address the potential clonality of *C. dubium* patches. Ramets were sampled from altogether 50 patches of *C. dubium* along the whole 400 km stretch of the Elbe River, where the species occurs (Figure 1, patch information in Supplement Table S1). 29 patches were located in the active floodplain and 21 in the inactive floodplain (behind the dykes without flooding). We used a regular grid design of 3 × 3 m with grid cells of 1 × 1 m resulting in 16 grid points. In each *C. dubium* patch 13 to 16 ramets were sampled. The patches found were often small and isolated without other patches in closer vicinity, i.e. the patch was often not much larger than the 3 × 3 m grid placed within. Other patches were embedded in larger stands. In September 2013, in 8 of the 50 patches, which were of larger size than the average patch, we sampled additional leaf material in a grid of 10 × 10 m using grid cells of 2 × 2 m, which did not overlap with the small grid. From the maximum of 36 samples per grid, 20-21 samples were randomly chosen for analysis. The size of each sampled *C. dubium* patch was assessed by inspecting the patch size itself and the surrounding area (including the grassland where the patch occurred and grasslands in closer vicinity). Small scale distribution maps available for the Elbe floodplain region of Lower Saxony and Saxony-Anhalt and local botanists were consulted as well. In sum, we derived a classification criterion for “patch size” as an important response variable for statistical analyses. Thereafter, 31 of the 50 patches were classified as “small” and 19 patches as “large” (information about patch properties: Supplement S1).

**Figure 1:**
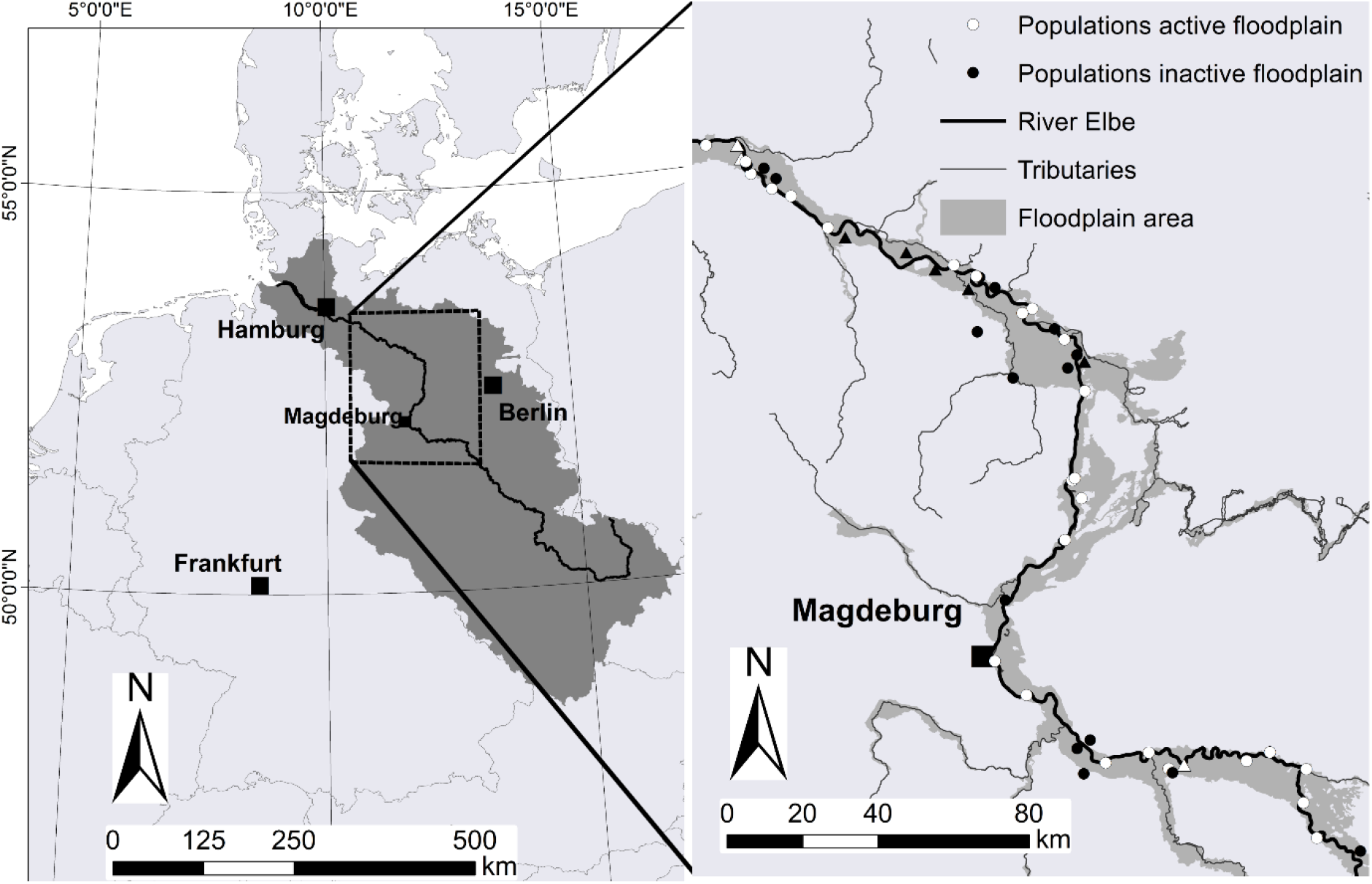
left) Elbe River catchment (dark grey) with the sampling area (outlined square); right) Spatial distribution of the 50 sampled *C. dubium*-patches along the Elbe River stretch where *C. dubium* can be found. Circles: location of sampling in 3 × 3 m grids; triangles: location of sampling in both 3 × 3 m and 10 × 10 m grids.

### Microsatellite and chloroplast DNA analysis

DNA was extracted from a total of 935 plants following the protocol as described in Dumolin et al. (1995) but using alkyltrimethylammonium bromide instead of cetyltrimethylammonium bromide and using 50µl of 1 × TE buffer with 10 µg/ml RNAse to resuspend the purified DNA. Genotyping of all samples was conducted using six nuclear microsatellite markers (nSSRs): CnD613, CnD806, CnD722, CnD723, CnD814, CnD817 (Michalczyk et al. 2012). PCR reactions were performed in a volume of 16.6 µl containing PCR-buffer and 0.65 units of Taq-polymerase (Molegene, Sinn, Germany), 0.3 mM dNTPs (Bioline, Luckenwalde, Germany), 16 mg/ml BSA (Thermo Fisher, St. Leon Rot, Germany), MgCl_2_ (Molegene) according to Table 1 and 20 ng of template DNA. Two different PCR profiles were used. For locus CnD723 a touchdown protocol was applied with an initial denaturation at 94 °C for 5 min followed by 10 cycles of 94 °C for 40 s, annealing at initially 59 °C – 1 °C after each cycle for 45 s and elongation at 72 °C for 40 s. Thereafter 20 cycles with a constant annealing temperature of 54 °C were performed followed by a final elongation at 72 °C for 10 min. The remaining microsatellite loci were amplified with the following protocol with individual annealing temperatures and hold times for denaturation and elongation according to Table 1: Initial denaturation at 94 °C for 5 min was followed by 30 cycles (for CnD613 35 cycles) of denaturation at 94 °C, annealing at the respective temperature for 45 s and elongation at 72 °C finalized by elongation at 72 °C for 10 min.

**Table 1:** Locus specific concentrations of MgCl2, annealing temperatures and hold times during PCR cycles for 6 nuclear microsatellite loci of *C. dubium*

| Locus | MgCl<sub>2</sub> conc. (mM) | Annealing temp. (°C) | Hold time (s) at 94°C/72°C |
| --- | --- | --- | --- |
| CnD814 | 3.0 | 51 | 45 |
| CnD722 | 3.0 | 56 | 40 |
| CnD806 | 3.0 | 56 | 30 |
| CnD613 | 3.0 | 54 | 45 |
| CnD817 | 2.4 | 59 | 30 |
| CnD723 | 3.0 | Touchdown, 54 | 40 |

The amplification products were separated by capillary electrophoresis using a MegaBACE 1000 automated sequencer (GE Healthcare, Freiburg, Germany). Fragment sizes were determined using the internal size standard MegaBACE ET400-R (60–400 bp; GE Healthcare) and alleles were scored with the software Fragment Profiler 1.2 (GE Healthcare).

For the analysis of chloroplast DNA (cpDNA) variation, we tested several regions of the chloroplast genome for sufficient polymorphism using universal primers. The most polymorphic regions, namely atpH/atpI (FM), trnT/trnF (TF) (Grivet et al. 2001) and trnH/trnK (HK) (Demesure et al. 1995) were used as markers in this study. One representative of each genet was analysed with the cpDNA markers and all samples which differed by just one allele from other genotypes; altogether 149 samples.

PCR reactions were performed in a volume of 30 µl with concentrations of Taq (Dream Taq, green, Thermo Fisher), dNTPs and BSA identical to the PCR protocol above. Concentrations of MgCl2 were 2 mM for FM and HK and 2,63 mM for TF. An amount of 35 ng of template DNA was added to each reaction. The temperature profiles were initial denaturation at 94 °C for 5 min, 40 (FM, HK) respectively 45 (TF) cycles of denaturation at 94 °C for 1 min, annealing at 56 °C (FM), 62 °C (HK) for 1 min or at 55 °C for 1 min 40s (TF), elongation at 72 °C for 1 min 30s (FM) or 1 min 40 s (HK, TF). All profiles were finalized with an elongation step at 72 °C for 10 min. PCR products were sent for clean-up and sequencing to the company LGC Genomics (Berlin, Germany). Sequences were analysed with the software CodonCode Aligner Version 7.1.2. From the sequence data, multilocus chloroplast haplotypes (hereinafter just called “haplotypes”) were identified.

Although chloroplast DNA is known to be maternally inherited in many angiosperms, this has not been determined specifically for *C. dubium* so far. To be on the safe side, we analysed multilocus haplotypes with the above mentioned cpDNA markers for four mother plants from the Elbe River from which we also collected seeds for cultivating off-spring. In all cases the chloroplast haplotype of the seedlings corresponded with that of the mother plants which suggests maternal inheritance of cpDNA in *C. dubium*.

### Handling of samples with presumed somatic mutations

In 16 out of the 50 3 × 3 m grids and four out of the eight 10 × 10 m grids samples occurred, which differed by just one SSR allele from other samples of the same grid forming a clone group. In most cases the fragment length difference corresponded to one single repeat motif. These samples were analysed several times in order to exclude genotyping errors. However, the differences remained after this procedure. The most plausible explanation is that these differences are the result of somatic mutations, which are predicted to be associated with the accumulation of somatic mutations in cell lineages (Schultz & Scofield 2009). It is unlikely that these differences could be the result of sexual reproduction, which is emphasized by the fact that we did not find samples with these small allele differences with differing chloroplast haplotypes. Since the assumed mutation model for microsatellites is a stepwise mutation of repeat motifs, we used a conservative approach and only assumed somatic mutation for those genotypes that differed by one allele only and therein by one repeat motif only (Ohta & Kimura 1973). Consequently, we assigned these peculiar samples to the respective genet for further analyses (see genotype table, Supplement S2).

### Data analysis

For the identification of clonal structures we used the “Multilocus Matches” option as implemented in GenAlEx 6.503 (Peakall and Smouse, 2012). To assess the power of the multilocus system to differentiate genotypes, the probability of identity for unrelated individuals (PI) and siblings (PI_sibs_) was estimated using GenAlEx 6.503. To estimate intra-patch clonal diversity, we determined clonal diversity R as R = (G - 1)/(n - 1), where G is the number of genets and n is the number of sampled ramets (Dorken & Eckert 2001).

In order to analyse the effects of patch and floodplain parameters on the spatial organisation of genets and ramets we screened each pair of samples within the grids for the presence of different genets. Based on this we introduced as response variable “probability of occurrence of non-clonal ramet pairs”. The presence/absence of non-clonal ramet pairs (ramet pairs of different genets) within all sampled grids together was related to small scale geographic distance using generalised linear models (GLM, with a binomial error structure). The geographic distances ranged from 1 - 4.24 m in the case of the 3 × 3 m grids and 2 - 14.14 m in the case of the 10 × 10 m grids. Further explanatory variables that were taken into account included the location of the patch relative to the dyke (active and inactive floodplain, variable “floodplain type”), the patch size in which the grid was embedded (large/small) and the number of chloroplast haplotypes in each studied patch. Furthermore, we calculated the genetic distance for each non-clonal ramet pair as response variable according to Smouse & Peakall (1999) using GenAlEx 6.503. For this, pairwise genetic distances of zero (two ramets of the same genet) were removed from each grid. Samples with somatic mutations assigned to clone groups were excluded as well. Genetic distances were then analysed in relation to the above mentioned variables “geographic distance”, “floodplain type”, “patch size” and “haplotype number” using GLM with a poisson distributed error structure.

Clonal diversity of the patches was related to several explanatory variables regarding properties of the patches (patch size, haplotype number) and location (floodplain type: active/inactive; distance to river, location along the river: middle/lower stretch) using analysis of variance and linear regression. Models were checked for homoscedasticity and normal distribution of errors using diagnostic plots. The analyses were performed using R 3.4.2. (R Development Core Team, 2017). Further analyses of genetic structure and genetic diversity were based on a reduced dataset including only a single representative of each clone (in total 121 individuals). We performed a Bayesian clustering analysis using STRUCTURE 2.3.4 (Pritchard et al. 2000) to search for evidence of population genetic structure. We applied STRUCTURE using the default settings (admixture model, correlated allele frequency model) and simulating for k= 1 to 8 with 5 replications each with 100,000 MCMC steps for burnin and 100,000 steps after burnin. Results were evaluated using the CLUMPAK pipeline (Kopelman et al. 2015) for visualization of the barplots and for applying the delta K method according to Evanno et al. (2005). Furthermore, a Principal Coordinates Analysis (PCoA) based on pairwise genetic distances between genets as well as analyses of molecular variance (AMOVA) with 999 permutations were performed in order to detect differentiation between patches of the active and inactive floodplain as well between the middle and the lower stretch of the Elbe river. Standard measures of genetic differentiation such as *G’*_*st*_ (Hedrick, 2005) and *D*_*est*_ (Jost, 2008) were determined among the same four groups of patches. These analyses were performed with the software GenAlEx 6.503.

## Results

The probability of identity (PID) of two randomly drawn individuals exhibiting the same genotype was 2.3 × 10^−7^ (1:4.4 million), for siblings 4.4 × 10^−3^ (1:226). This low value of PID assured sufficient power of the six microsatellite markers to differentiate among genets (Waits et al. 2001). From the altogether 935 analysed samples in the 3 × 3 m and 10 × 10 m grids, 121 different genotypes could be detected after removing genotypes with putative somatic mutations. From the 50 patches sampled in the 3 × 3 m grid, 31 patches exhibited only one single genet each forming a large ramet group. The 19 other patches (13 belonging to large sized, 6 to small patches contained 2 to 4 different genets each, but 4-genet patches were restricted to large patches of the inactive floodplain. Patches sampled in 10 × 10 m grids exhibited 2 to 8 different genotypes each (mean genets per patch in the active/inactive floodplain: 2.7/6.4). Ramets of the same genet could be detected over a distance of more than 14 metres (diagonal edges of the large grid). A genet never occurred in more than one patch. This indicates that vegetative dispersal units were not dispersed over large distances between the investigated locations.

Altogether 18 multilocus chloroplast haplotypes could be detected of which 4 occurred frequently (frequency > 10%). Within each of 50 patches sampled with the 3 × 3 m grid one (40 patches) or two haplotypes (10 patches) occurred. In each of six patches sampled with the 10 × 10 m grid also one (3 patches) or two haplotypes (3 patches) occurred, while in two single patches three and four haplotypes were detected.

### Small scale: genetic patterns within patches

The probability of occurrence of non-clonal ramet pairs (=ramet pairs of different genets) was significantly related to the geographic distance in both the 3 × 3 m and 10 × 10 m grid with an increase in probability of occurrence with increasing distance (Figure 2, Table 2). Additionally, the occurrence of non-clonal ramet pairs was more likely in patches of larger size and in the inactive floodplain. Also, it was more likely in patches exhibiting two instead of one haplotype (3 × 3 m grids) (Figs. 2 a,b). In the case of the 10 × 10 m grids the probability of occurrence increased from 1 to 4 haplotypes present in the grids (Figure 2 d).

**Figure 2:**
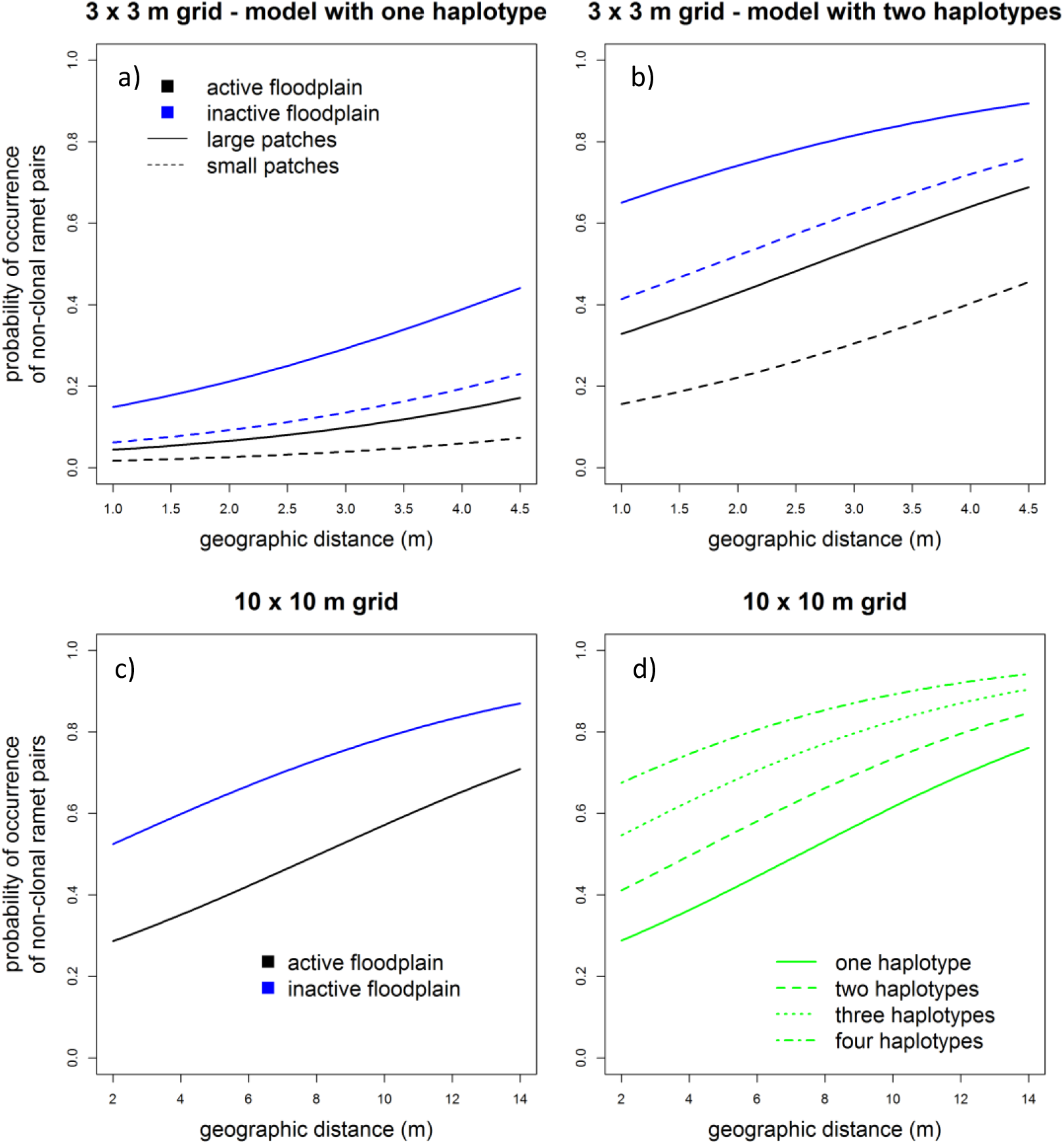
Probability of occurrence of non-clonal ramet pairs in relation to “geographic distance” and “floodplain type” in a) 3 × 3 m grids with one haplotype present, b) 3 × 3 m grids with two haplotypes present (here the “patch size” also included), c) 10 × 10 m grids and d) 10 × 10 m grids in relation to the number of haplotypes present in the grids.

**Table 2:** Results of GLM (binomial-family) with presence/absence of non-clonal ramet pairs as response variable. Data set: Samples of the 3 × 3 m grid from 50 patches and samples of the 10 × 10 m grid from 8 patches. Explained deviance 3 × 3 m grid = 24.1 %; 10 × 10 m grid = 10.1 %. Bonferroni adjusted alpha levels: small grid: 1.78 × 10^−7^ (0.001/5598), large grid: 6.49 × 10^−7^ (0.001/1540).

|  | Estimate | Std. Error | z-value | p |
| --- | --- | --- | --- | --- |
| **3 x 3 m grid** |  |  |  |  |
| Intercept | -4.546 | 0.182 | -24.941 | < 0.001 *** |
| Geographic distance | 0.431 | 0.049 | 8.811 | < 0.001 *** |
| Patch size | -0.970 | 0.086 | -11.275 | < 0.001 *** |
| Floodplain type | -1.338 | 0.093 | -14.442 | < 0.001 *** |
| Haplotype number | 2.367 | 0.095 | 24.945 | < 0.001 *** |
| **10 x 10 m grid** |  |  |  |  |
| Intercept | -1.240 | 0.199 | -6.234 | < 0.001 *** |
| Geographic distance | 0.161 | 0.021 | 7.513 | < 0.001 *** |
| Floodplain type | -0.819 | 0.114 | -7.213 | < 0.001 *** |
| Haplotype number | 0.465 | 0.062 | 7.455 | < 0.001 *** |

Genetic distances among non-clonal ramet pairs in the 3 × 3m grids were significantly higher in patches of larger size and where two different haplotypes were present (Figure 3a,b, Table 3). In the 10 × 10 m grids, genetic distance was higher in the inactive floodplain and increased with the number of haplotypes present. “Geographic distance” was not related to genetic distance in both grid sizes (Figs. 3c,d, Table 3).

**Figure 3:**
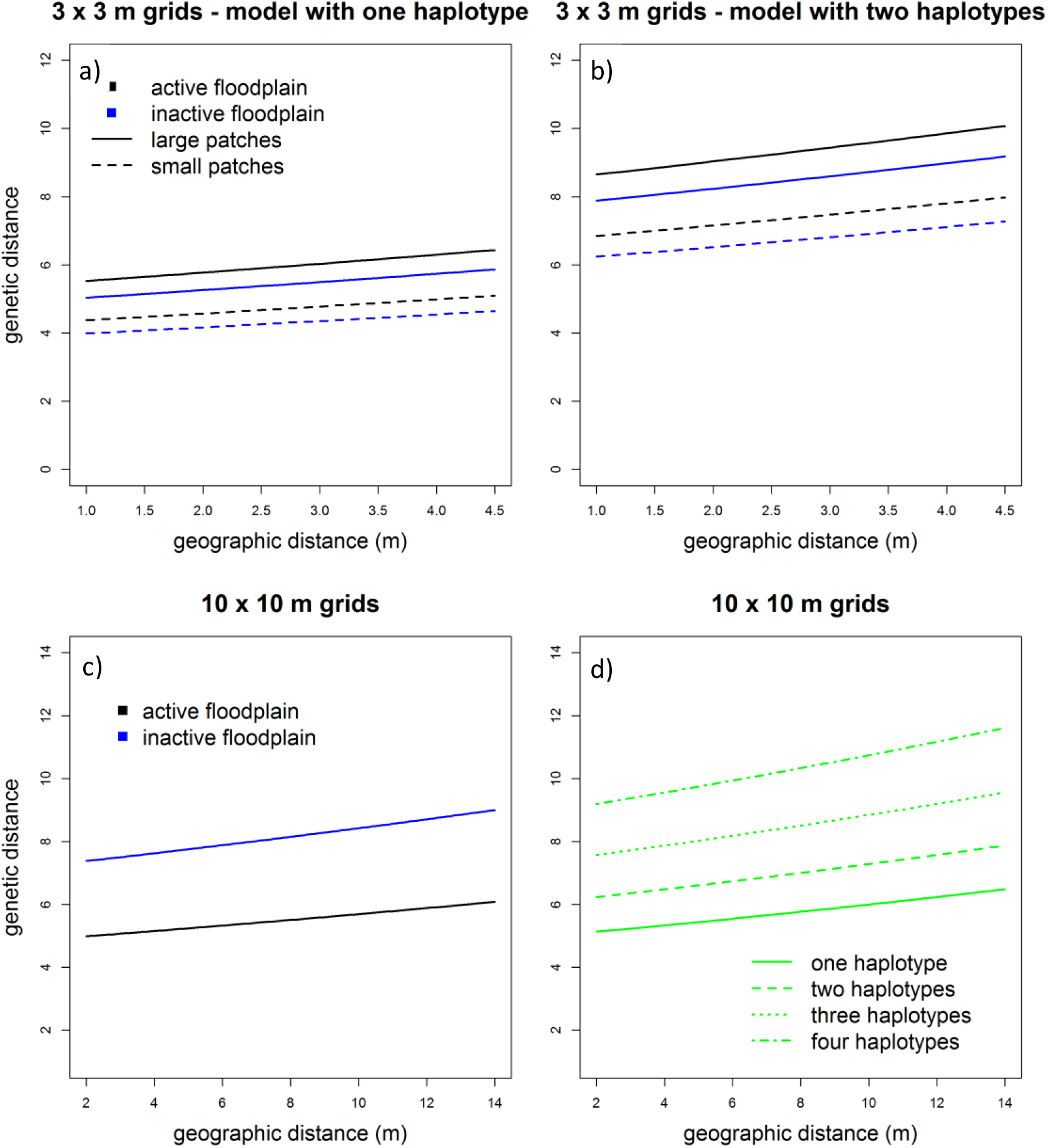
a) Genetic distances of non-clonal ramet pairs (ramet pairs of same genets excluded) in relation to “geographic distance”, “floodplain type” and “patch size” in 3 × 3 m grids with one haplotype present and b) with two haplotypes present, c) Genetic distances of non-clonal ramet pairs in relation to “geographic distance” and “floodplain type” and d) in relation to “geographic distance” and “number of haplotypes” in 10 × 10 m grids.

**Figure 4:**
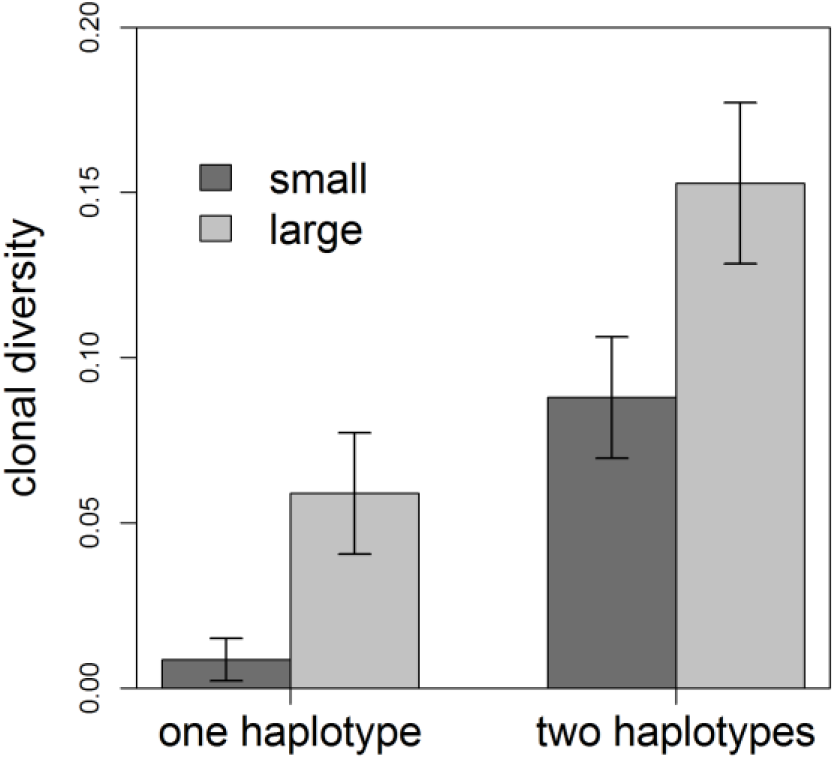
Intra-patch clonal diversity in relation to size and haplotype number of the patch. Due to the small number of patches sampled in 10 × 10 m grids only the 50 3 × 3 m grids were considered.

**Table 3:** Results of GLM (poisson-family) with genetic distance as response variable. Data set: Samples of the 3 × 3 m grid from 50 patches and samples of the 10 × 10 m grid from eight patches. Explained deviance 3 × 3 m grid = 22.1 %; 10 × 10 m grid = 19.3 %. Bonferroni adjusted alpha levels: 3 × 3 m grid: 1.12 × 10^−6^ (0.001/888), 10 × 10 m grid: 1.13 × 10^−6^ (0.001/879)

|  | Estimate | Std. Error | z-value | p |
| --- | --- | --- | --- | --- |
| **3 x 3 m grid** |  |  |  |  |
| Intercept) | 1.124 | 0.059 | 18.959 | < 0.001 *** |
| Geographic distance | 0.044 | 0.015 | 2.894 | ns |
| Patch size | -0.233 | 0.029 | -8.139 | < 0.001 *** |
| Floodplain type | 0.093 | 0.029 | 3.267 | ns |
| Haplotype number | 0.448 | 0.029 | 15.540 | < 0.001 *** |
| **10 x 10 m grid** |  |  |  |  |
| (Intercept) | 1.555 | 0.047 | 33.341 | < 0.001 *** |
| Geographic distance | 0.017 | 0.005 | 3.808 | ns |
| Floodplain type | -0.246 | 0.033 | -7.361 | < 0.001 *** |
| Haplotype number | 0.156 | 0.012 | 13.228 | < 0.001 *** |

### Large scale: genetic patterns along the Elbe River

Considering the intra-patch clonal diversity (based on the 50 3 × 3 m grids) it was significantly related to the patch properties “patch size” and “haplotype number” (Table 4) while variables related to the spatial distribution of patches (floodplain type, distance to river, river stretch, river kilometre) had no significant effect.

**Table 4:** Result of analysis of variance with intra-patch clonal diversity in relation to patch size and number of haplotypes per patch.

|  | **d.f.** | **MS** | **F-value** | **p** |
| --- | --- | --- | --- | --- |
| **Patch size** | 1 | 0.04296 | 21.75 | <0.001 *** |
| **Haplotype number** | 1 | 0.054 | 23.79 | <0.001 *** |
| **Residuals** | 47 | 0.00227 |  |  |

Regarding the genetic diversity and differentiation parameters no differences between recent and older floodplain and middle and lower stretch of the Elbe River could be detected (Table 5). The PCoA showed no pattern for the grouping of samples neither for the river course nor for the floodplain type. This is consistent with the results of the Bayesian cluster analysis, which did not provide any evidence for population structure (Supplement S3).

**Table 5:** Average values of genetic diversity parameters (number of alleles: N_a_; effective number of alleles: N_e_; observed (H_o_) and expected (H_e_) heterozygosity; inbreeding coefficient: (F) for the active versus inactive floodplain patches as well as for the patches of the middle stretch of the Elbe River versus lower stretch patches.

| **Pop** | **$N_a$** | **$N_e$** | **$H_o$** | **$H_e$** | **F** |
| --- | --- | --- | --- | --- | --- |
| **active** | 11.333 | 5.524 | 0.641 | 0.735 | 0.117 |
| **inactive** | 10.000 | 4.979 | 0.655 | 0.726 | 0.090 |
| **middle** | 9.833 | 5.401 | 0.653 | 0.753 | 0.114 |
| **lower** | 11.000 | 5.137 | 0.647 | 0.719 | 0.095 |

The results of respective AMOVAs indicated that only 1-2 % of the molecular variance resided among populations (i.e. among active and inactive floodplain, among middle and lower stretch). Differentiation parameters *G’*_*st*_ and *D*_*est*_ values were also very low (active vs. inactive floodplain: 0.019/0.016; middle vs. lower stretch: 0.024/0.020).

## Discussion

The analyses of the population genetic structure in *C. dubium* along the Elbe River revealed clear results on both considered spatial scales. Within-patch clonal diversity was found to be extremely low. Clear patterns were evident regarding the spatial organisation of clonal structures and their occurrence in relation to patch size and floodplain type. In contrast, we observed a complete lack of genetic structure on the large scale i.e. that neither differences between active and inactive floodplain nor along the river course regarding genetic diversity and differentiation parameters could be found.

Microsatellite marker analysis revealed that almost two thirds of the studied grids consisted of only one genet exhibiting a guerrilla like growth form (Lovett-Doust 1981). The results show clearly, that especially the small patches, not surrounded by other patches in the vicinity, are often monoclonal. It can be assumed that isolation of these monoclonal ramet groups provoke a lack of gene flow via pollen between genets which may be a reason for the absence of sexual reproduction. Furthermore, there is growing evidence that clonal growth can have negative effects on sexual reproduction e.g. due to strong trade-offs between investment in sexual and vegetative reproduction leading to limited allocation to flowering and seed production in case of rapid clonal expansion. Another aspect deals with the absence of mating groups required for outcrossing in large monoclonal ramet groups leading to inbreeding depression and pollen discounting associated with geitonogamous self-pollination (Barrett 2015). In the case of *C. dubium* a mixed mating system with cross- and self-fertilisation is mentioned in the literature (Chytrý et al. 2021). The observed inbreeding coefficients might indicate a moderate level of inbreeding. However, we may have introduced a bias by constructing “artificial populations” following our spatial categories of middle/lower river stretch or active/inactive floodplain for the purpose of estimating this coefficient. On the other hand, the observed levels of heterozygosity do not show evidence for high rates of self-fertilisation. Therefore, we think that the evidence rather supports a predominantly outcrossing mating system in a population not in equilibrium due to high levels of clonal reproduction (Reichel et al. 2016) and genetic isolation of the patches. This could result in the lack of seed development or viable seeds if mating partners are not present. Indeed, a low germination rate was shown in several germination experiments with *C. dubium* seeds of Rhine and Elbe River populations. Germination percentage turned out to be much lower than in other floodplain- and flood-meadow species (Hölzel & Otte 2004, Hölzel 2005, Geissler & Gzik 2008, 2010).

Furthermore, clonal plants have indeterminate growth and are potentially immortal. The potentially high number of mitotic divisions between zygote formation and gamete production is considered to be associated with the accumulation of somatic mutations leading to slight DNA variation within clonal lineages. They are increasingly recognised as an important feature in clonally reproducing plants and long-lived trees (Schultz & Scofield 2009, Barrett 2015). Somatic mutations at microsatellite loci have been observed e.g. in Rosaceae as well as Salicaceae species (see Jankowska-Wroblewska et al. 2016 for a summarisation). Indeed, also in our study samples occurred with single allele differences compared to adjacent groups of otherwise identical genotypes, which we considered to be somatic mutations. This points toward a comparatively high age of the corresponding genets (Ally et al. 2008). Somatic mutations are seen as a cause for the reduction in fertility in clonal plants due to a gradual accumulation of deleterious mutations transmitted from shoot meristems to gametes (Barrett 2015). This could be a further reason for the observed low clonal diversity in studied *C. dubium* patches. Taken the facts as a whole, it can be assumed that the observed prolonged clonal growth may be a successful strategy to secure population persistence after the loss in area, connectivity and quality of habitats in the short term (de Witte & Stöcklin 2010). In the long term, however, this strategy might be detrimental.

By including the results of the cpDNA marker analysis, additional information on gene flow processes can be derived (Ziegenhagen et al. 2003). In about half of the patches, where more than one genet occurred, two haplotypes could be found. This is evidence that seeds of different mother plants germinated when these patches were established. It is not completely excluded in the one-haplotype patches that more than several mother plants with the same haplotype contributed seeds to the patch. However, it is more likely, that in most cases pollen from surrounding genets contributed to gene flow and that the different genets of the patch consisted of relatives thus having the same haplotype. This assumption is strongly supported by the higher genetic distances between ramets in two-haplotype patches in comparison with those in one-haplotype patches. We can conclude that gene flow by pollen and seeds as well as sexual reproduction is possible but it is strongly associated with large patches.

Surprisingly, the results indicate that sexual reproduction is at least slightly enhanced in the inactive compared to the active floodplain which is especially pronounced in the 10 × 10 m grid plots. Here, the number of genets per plot in the inactive floodplain was more than twice as high as in the active floodplain. Generally, reasons for the lack of sexual reproduction have often been associated with the lack of bare ground which is seen as a prerequisite for germination in many herbal plants and which is suggested for *C. dubium* as well (Geissler & Gzik 2010). If this is a main factor triggering sexual reproduction, a higher clonal diversity would be assumed in the active floodplain since erosional and depositional processes due to river flow are present and even accelerated in most cases due to the pronounced narrowing of the floodplains by dykes. However, the contrary is true in our study. Mosner et al. (2012) found also a higher clonal diversity in *Salix viminalis* stands in the inactive floodplain of the Elbe River. They identified mechanical forces through river flow in combination with long time flooding as drivers for intensive vegetative resprouting in the active floodplain. For *C. dubium*, Geissler & Gzik (2008) stated negative effects of extended flooding on seeds through inhibition of germination as well as remarkable decay rates which could be of higher importance in the active floodplain.

Regarding the genetic structure at the large scale considering the whole river stretch a clear lack of a genetic pattern can be stated. This finding suggests at first glance that gene flow acts efficiently across large distances along the Elbe River and also across the different floodplain types (active and inactive). It seems that one continuous population is present at the Elbe River which would be in line with an observed low differentiation of plant populations in dynamic river systems due to hydrochory (Hu et al. 2010). However, the above explained clonal organization of *C. dubium* within the studied patches provided a contrasting picture. The results suggest that the large scale lack of population structure observed in *C. dubium* along the Elbe River might not be the result of extensive gene flow across the landscape. It is justified to assume that gene flow is extremely low caused by the strong isolation of patches which hampers pollination. Moreover, gene flow by seeds seems to be even more restricted than by pollen due to limited germination and seedling establishment. Grassland intensification or abandonment in the active and inactive floodplain as well as the conversion into arable land in the inactive floodplain together with hydrological and hydraulic alterations of the river systems are the main reasons for the decline of the *Cnidium* flood-meadows during the last decades (Finck et al. 2017) and therefore the reason for the assumed isolation of *C. dubium* patches in the river landscape.

Many patches may be remnants of former times in which *C. dubium* was more abundant due to environmental conditions more suitable for this plant. It seems to be conclusive that *C. dubium* is representing somewhat like a frozen population in the Elbe River region. This might explain the observed elevated inbreeding coefficients to some extent, since these might just reflect a population not in equilibrium. Thus *C. dubium* may be predestined to further decline due to the particularly low number of genets, their isolation through habitat fragmentation and deterioration, as well as restricted sexual reproduction. There is reason to presume that *C. dubium* is in the process of time-delayed extinction even without any further habitat loss occurring. Thus an extinction debt for the Elbe river population can be stated.

Considerable efforts are made by nature conservation authorities e.g. in the Elbe River and Upper Rhine River region to restore *Cnidium* flood-meadows. Hay transfer and application of threshing material from species-rich source stands containing seeds from target species proved to be a successful approach for restoration (Kiehl et al. 2010, Bischoff et al. 2018) while it is insufficient only to apply nature conservation management on degraded floodplain grasslands due to strong dispersal limitations of the target species (Donath et al. 2003) which is surely true for *C. dubium* as well. However, while many of these species arise after hay transfer, this measure is obviously not very successful in the case of *C. dubium* due to low germination rates of seeds transferred from the source populations (Hölzel et al. 2006). Instead, ramets of a number of different genets with parts of their roots should be planted in groups, which is easily possible as we could observe in a small side experiment within this study. We cut out three to four connected non-flowering ramets (including root and some soil) of eight studied patches with subsequent successful establishment in a common flowerbed. In the subsequent year many of the ramets of the verifiably different genets flowered and developed seeds. The subsequent sowing of seeds resulted in high germination rates of more than 70% (unpublished data). The open flowering of ramet groups of different genets obviously enhanced outcrossing and therefore successful sexual reproduction. Since no genetic differentiation could be determined the introduction of genets from a number of different patches via parts of roots seems promising for *C. dubium* re-establishment in restored flood meadows.

## Conclusions

Having in mind the strong decline of this species in Central Europe, the findings give reason for great concern. The effective population size is much lower than the number of plants suggests and due to the identified extinction debt we can assume that the species is doomed to become extinct at the Elbe River. However, the identification of an unpaid extinction debt implies that there still is a chance to counteract the future extinction by targeted habitat restoration and conservation actions (Kuussaari et al. 2009). Nature conservation agencies should seize this opportunity and include suitable re-introduction measures into floodplain meadow restoration and conservation programs to keep *C. dubium* populations alive.

## Acknowledgement

We thank Horst Jage (botanist, Kemberg) and Christiane Schreck (administration of the Biosphere Reserve “Elbe River Landscape”, Lower Saxony, Hitzacker) for giving valuable information on *C. dubium* locations at the Elbe River region. Furthermore, we are grateful to Lisa Thomas and to Filine Seele for sampling support and laboratory work. The authors declare no conflicts of interest. The study was funded by the German Research Foundation (DFG grant LE 1364/5-1).

## Author’s contributions

IL, SL and BZ conceived the ideas and designed methodology; IL, EM and CM collected the data; IL, SL and EM analysed the data; IL led the writing of the manuscript. All authors contributed critically to the drafts and gave final approval for publication.

## Data archiving statement

We intend to archive genotyping data of all collected samples for the microsatellite as well as cpDNA data in the Dryad Digital Repository (http://datadryad.org/)

## Notes

### Competing Interest Statement

The authors have declared no competing interest.

