## Supplement S1 for "*Predominance of clonal propagation* conceals extinction risks of the highly endangered floodplain herb *Cnidium dubium*"

### Supplement Table S1
### Patch information of the 50 patches sampled in 3x3m grids and of the 8 patches sampled in 10x10m grids

| Number | Label | Location name | Size | Number genets | Number haplotypes | Floodplain type | N | Location along the Elbe River | River kilometre | Xkoord |
|---|---|---|---|---|---|---|---|---|---|---|
| **Samples of the 3x3 m grid** | | | | | | | | | | |
| 1 | HT | Hittbergen | small | 1 | 1 | active | 16 | lower | 566.9 | 3206470 |
| 2 | BR | Brackede | large | 1 | 1 | active | 16 | lower | 557.4 | 3214756 |
| 3 | HB | Heisterbusch | large | 3 | 2 | active | 15 | lower | 552.4 | 3215454 |
| 4 | BL | östl. Bleckede | small | 1 | 1 | active | 16 | lower | 547.8 | 3217677 |
| 5 | WH | Neu Wendischthun | large | 4 | 2 | inactive | 16 | lower | 550.8 | 3216870 |
| 6 | WT | Neu Wendischthun | small | 3 | 2 | active | 15 | lower | 552.3 | 3216567 |
| 7 | SP | östl. Stiepelse | large | 1 | 1 | inactive | 16 | lower | 545.4 | 3221254 |
| 8 | NG | nördl. Gülstorf | large | 2 | 1 | inactive | 16 | lower | 539.9 | 3224226 |
| 9 | GK | Groß Kühren | large | 2 | 1 | active | 14 | lower | 534.6 | 3227783 |
| 10 | WG | Walmsburg | small | 1 | 1 | active | 16 | lower | 540.0 | 3222868 |
| 11 | SE | bei Seedorf | large | 2 | 1 | inactive | 15 | lower | 517.9 | 3241180 |
| 12 | LA | Laake | large | 1 | 1 | active | 15 | lower | 521.7 | 3236806 |
| 13 | BZ | Breetz | large | 4 | 1 | inactive | 16 | lower | 488.6 | 3256988 |
| 14 | RE | Restorf | large | 4 | 1 | inactive | 16 | lower | 480.6 | 3264253 |
| 15 | WZ | Wanzer | large | 1 | 1 | inactive | 16 | lower | 465.9 | 3272570 |
| 16 | LB | Lindenberg | small | 1 | 1 | inactive | 16 | lower | 458.8 | 3273985 |
| 17 | WW | westl. Wittenberge | small | 2 | 2 | inactive | 16 | lower | 456.2 | 3279615 |
| 18 | WD | westl. Wentdorf | small | 1 | 1 | inactive | 16 | lower | 468.8 | 3275794 |
| 19 | LE | Lennewitzer Eichen | small | 1 | 1 | inactive | 16 | lower | 432.9 | 3294441 |
| 20 | TO | NSG Tonabgrabungen | large | 4 | 2 | inactive | 16 | lower | 420.9 | 3301573 |
| 21 | WU | Wulkau | small | 1 | 1 | active | 16 | lower | 411.5 | 3301019 |
| 22 | KA | Kannenberg | small | 1 | 1 | inactive | 16 | lower | 418.3 | 3297044 |
| 23 | RÄ | Räbel | small | 1 | 1 | inactive | 15 | lower | 421.2 | 3299743 |
| 24 | OS | Osterburg | small | 1 | 1 | inactive | 11 | lower | 440.7 | 3282402 |
| 25 | CU | Cumlosen | small | 1 | 1 | active | 16 | lower | 468.5 | 3274890 |
| 26 | LN | Lütkenwisch Nord | small | 1 | 1 | active | 16 | lower | 476.4 | 3269275 |
| 27 | BD | Bälow Deich | large | 2 | 1 | active | 16 | lower | 445.7 | 3288986 |

| | | | | | | | | | | |
|---|---|---|---|---|---|---|---|---|---|---|
| 28 | WE | Werben | small | 1 | 1 | active | 16 | lower | 429.5 | 3296778 |
| 29 | BA | Beuster Altarm | small | 1 | 1 | active | 16 | lower | 443.5 | 3286287 |
| 30 | FA | Fischbeck Altarm | small | 1 | 1 | active | 14 | middle | 387.5 | 3295712 |
| 31 | FD | Fischbeck Deich | large | 1 | 1 | active | 16 | middle | 387.9 | 3296482 |
| 32 | BB | Bucher Brack | large | 3 | 2 | active | 14 | middle | 381.3 | 3297789 |
| 33 | PA | Parey | small | 1 | 1 | active | 15 | middle | 369.5 | 3292542 |
| 34 | GH | Zw. Glindenberg und Heinrichsberg | small | 1 | 1 | inactive | 16 | middle | 341.9 | 3275474 |
| 35 | CR | Westl. Cracau Nähe Elbufer | small | 2 | 2 | active | 15 | middle | 323.0 | 3271353 |
| 36 | SBE | Vorland östl. Schönebeck | small | 1 | 1 | active | 16 | middle | 309.2 | 3278950 |
| 37 | BH | südl. Breitenhagen | large | 2 | 1 | inactive | 16 | middle | 283.5 | 3291154 |
| 38 | WB | Wulfener Bruch | large | 1 | 1 | inactive | 16 | middle | 277.3 | 3292361 |
| 39 | AK | nordöstl. Aken | small | 2 | 2 | active | 15 | middle | 274.0 | 3298340 |
| 40 | SW | Wiesen am Schwedenhaus | small | 1 | 1 | active | 16 | middle | 245.6 | 3314857 |
| 41 | RL | Südl. Rosslau | small | 1 | 1 | active | 16 | middle | 257.6 | 3309985 |
| 42 | ST | bei Steckby | small | 1 | 1 | inactive | 16 | middle | 282.0 | 3294778 |
| 43 | SB | direkt unter Stromleitung Biores | small | 1 | 1 | inactive | 16 | middle | 245.3 | 3315835 |
| 44 | VO | östl. Vockerode | large | 2 | 1 | active | 14 | middle | 244.4 | 3318989 |
| 45 | WiB | gegenüber Wittenberg | small | 3 | 1 | active | 14 | middle | 217.9 | 3335585 |
| 46 | WL | östl. Wittenberg/Lutherbrunnen | small | 1 | 1 | active | 15 | middle | 210.0 | 3341921 |
| 47 | EL | östl. Elster | large | 2 | 2 | active | 16 | middle | 199.1 | 3351151 |
| 48 | KL | südlich Klöden | small | 1 | 1 | active | 16 | middle | 188.7 | 3349634 |
| 49 | PR | bei Proschwitz | small | 1 | 1 | active | 15 | middle | 176.5 | 3352460 |
| 50 | PD | NSG Prudel bei Dautzschen | small | 2 | 1 | inactive | 13 | middle | 162.8 | 3363676 |
| **Samples of the 10x10m grids** | | | | | | | | | | |
| 2b | NBR | Brackede großes Grid | large | 2 | 2 | active | 20 | lower | 557.4 | 3214756 |
| 3b | NHB | Heisterbusch großes Grid | large | 2 | 1 | active | 20 | lower | 552.4 | 3215454 |
| 11b | NSE | bei Seedorf großes Grid | large | 7 | 4 | inactive | 20 | lower | 517.9 | 3241180 |
| 13b | NBZ | Breetz großes Grid | large | 7 | 2 | inactive | 20 | lower | 488.6 | 3256988 |
| 14b | NRE | Restorf großes Grid | large | 7 | 1 | inactive | 20 | lower | 480.6 | 3264253 |
| 15b | NWZ | Wanzer großes Grid | large | 8 | 3 | inactive | 20 | lower | 465.9 | 3272570 |
| 20b | NTO | NSG Tonabgrabungen großes Grid | large | 3 | 1 | inactive | 20 | lower | 420.9 | 3301573 |
| 44b | NVO | östl. Vockerode großes Grid | large | 4 | 2 | active | 21 | middle | 244.4 | 3318989 |

| Ykoord |
| --- |
| 5921198 |
| 5920501 |
| 5916864 |
| 5912665 |
| 5915100 |
| 5915981 |
| 5913844 |
| 5910822 |
| 5905893 |
| 5908239 |
| 5893777 |
| 5896691 |
| 5888427 |
| 5883305 |
| 5877338 |
| 5865883 |
| 5877185 |
| 5880243 |
| 5864938 |
| 5855723 |
| 5847905 |
| 5854344 |
| 5857584 |
| 5852929 |
| 5880623 |
| 5884036 |
| 5870726 |

5861966
5869847
5824527
5825066
5819603
5808980

5794337

5778474
5768839
5753603
5746794
5749208
5746327
5751013
5755572

5745130

5746908
5746667

5748423

5743197
5734376
5724765
5720326

5920501
5916864
5893777
5888427
5883305
5877338
5855723
5746908
