## Supplement S2 for "*Predominance of clonal propagation* conceals extinction risks of the highly endangered floodplain herb *Cnidium dubium*"

### Supplement Table S2

### Genotype table for nuclear microsatellites of C. dubium patches containing somatic mutations.

Sample names are composed of a two or three letter identifier for the patch followed by the coordinates in the grid i.e. A-D (small grids) or A-F (large grids) for the X coordinate and 1-4 respectively 1-6 for the Y coordinate.

Cells filled red contain alleles, which are assumed to have arisen by somatic mutations. We assumed that the mutated alleles occured in the minority of ramets of the same clone. Differently colored lines within one patch indicate different genets

| Patch Information | Sample | Patch No. | Cnd806 | | CnD723 | | CnD722 | | CnD814 | | CnD613 | | CnD817 | |
|---|---|---|---|---|---|---|---|---|---|---|---|---|---|---|
| **Heisterbusch** | NHB-A1 | 333 | 113 | 113 | 107 | 107 | 85 | 87 | 203 | 203 | 140 | 142 | 125 | 125 |
| 10x10 m grid | NHB-A4 | 333 | 113 | 113 | 107 | 107 | 85 | 87 | 203 | 203 | 142 | 144 | 125 | 125 |
| | NHB-A6 | 333 | 113 | 113 | 107 | 107 | 85 | 87 | 203 | 203 | 142 | 144 | 125 | 125 |
| | NHB-B2 | 333 | 113 | 113 | 107 | 107 | 85 | 87 | 203 | 203 | 140 | 142 | 125 | 125 |
| | NHB-B4 | 333 | 113 | 113 | 107 | 107 | 85 | 87 | 203 | 203 | 142 | 144 | 125 | 125 |
| | NHB-B5 | 333 | 113 | 113 | 107 | 107 | 85 | 87 | 203 | 203 | 142 | 144 | 125 | 125 |
| | NHB-C2 | 333 | 113 | 113 | 107 | 107 | 85 | 87 | 203 | 203 | 142 | 144 | 125 | 125 |
| | NHB-C3 | 333 | 113 | 116 | 107 | 107 | 85 | 87 | 203 | 203 | 142 | 144 | 125 | 125 |
| | NHB-C4 | 333 | 113 | 113 | 107 | 107 | 85 | 87 | 203 | 203 | 140 | 142 | 125 | 125 |
| | NHB-C5 | 333 | 113 | 116 | 107 | 107 | 85 | 87 | 203 | 203 | 142 | 144 | 125 | 125 |
| | NHB-C6 | 333 | 113 | 116 | 107 | 107 | 85 | 87 | 203 | 203 | 142 | 144 | 125 | 125 |
| | NHB-D1 | 333 | 113 | 113 | 107 | 107 | 85 | 87 | 203 | 203 | 140 | 142 | 125 | 125 |
| | NHB-D3 | 333 | 113 | 113 | 107 | 107 | 85 | 87 | 203 | 203 | 142 | 144 | 125 | 125 |
| | NHB-D4 | 333 | 113 | 116 | 107 | 107 | 85 | 87 | 203 | 203 | 142 | 144 | 125 | 125 |
| | NHB-D6 | 333 | 113 | 116 | 107 | 107 | 85 | 87 | 203 | 203 | 142 | 144 | 125 | 125 |
| | NHB-E2 | 333 | 113 | 113 | 107 | 107 | 85 | 87 | 203 | 203 | 140 | 142 | 125 | 125 |
| | NHB-E3 | 333 | 113 | 113 | 107 | 107 | 85 | 87 | 203 | 203 | 140 | 142 | 125 | 125 |
| | NHB-E5 | 333 | 113 | 116 | 107 | 107 | 85 | 87 | 203 | 203 | 142 | 144 | 125 | 125 |
| | NHB-F1 | 333 | 113 | 113 | 107 | 107 | 85 | 87 | 203 | 203 | 140 | 142 | 125 | 125 |
| | NHB-F4 | 333 | 113 | 113 | 107 | 107 | 85 | 87 | 203 | 203 | 140 | 142 | 125 | 125 |
| **Neu Wendischthun** | WT-A1 | 6 | 101 | 113 | 105 | 109 | 89 | 89 | 191 | 227 | 94 | 94 | 125 | 125 |

| | | | | | | | | | | | | | | |
|---|---|---|---|---|---|---|---|---|---|---|---|---|---|---|
| 3x3 m grid | WT-A2 | 6 | 89 | 116 | 105 | 109 | 87 | 87 | 203 | 203 | 100 | 118 | 125 | 127 |
| | WT-A3 | 6 | 101 | 113 | 105 | 109 | 89 | 89 | 191 | 227 | 94 | 94 | 125 | 125 |
| | WT-A4 | 6 | 89 | 107 | 105 | 107 | 85 | 89 | 203 | 230 | 114 | 116 | 125 | 125 |
| | WT-B1 | 6 | 89 | 119 | 105 | 109 | 87 | 87 | 203 | 203 | 100 | 118 | 125 | 127 |
| | WT-B2 | 6 | 89 | 119 | 105 | 109 | 87 | 87 | 203 | 203 | 100 | 118 | 125 | 127 |
| | WT-B3 | 6 | 89 | 119 | 105 | 109 | 87 | 87 | 203 | 203 | 100 | 118 | 125 | 127 |
| | WT-C1 | 6 | 89 | 119 | 105 | 109 | 87 | 87 | 203 | 203 | 100 | 118 | 125 | 127 |
| | WT-C2 | 6 | 89 | 119 | 105 | 109 | 87 | 87 | 203 | 203 | 100 | 118 | 125 | 127 |
| | WT-C3 | 6 | 89 | 119 | 105 | 109 | 87 | 87 | 203 | 203 | 100 | 118 | 125 | 127 |
| | WT-C4 | 6 | 89 | 107 | 105 | 107 | 85 | 89 | 203 | 230 | 114 | 116 | 125 | 125 |
| | WT-D1 | 6 | 89 | 119 | 105 | 109 | 87 | 87 | 203 | 203 | 100 | 118 | 125 | 127 |
| | WT-D2 | 6 | 89 | 116 | 105 | 109 | 87 | 87 | 203 | 203 | 100 | 118 | 125 | 127 |
| | WT-D3 | 6 | 89 | 107 | 105 | 107 | 85 | 89 | 203 | 230 | 114 | 116 | 125 | 125 |
| | WT-D4 | 6 | 89 | 107 | 105 | 107 | 85 | 89 | 203 | 230 | 114 | 116 | 125 | 125 |
| **Groß Kühren** | GKA1 | 9 | 107 | 143 | 107 | 111 | 85 | 85 | 203 | 203 | 114 | 116 | 125 | 125 |
| 3x3 m grid | GKA2 | 9 | 107 | 143 | 107 | 111 | 85 | 85 | 203 | 203 | 114 | 116 | 125 | 125 |
| | GKA3 | 9 | 107 | 143 | 107 | 111 | 85 | 85 | 203 | 203 | 114 | 116 | 125 | 125 |
| | GKA4 | 9 | 107 | 143 | 107 | 111 | 85 | 85 | 203 | 203 | 114 | 118 | 125 | 125 |
| | GKB1 | 9 | 107 | 143 | 107 | 111 | 85 | 85 | 203 | 203 | 114 | 118 | 125 | 125 |
| | GKB2 | 9 | 107 | 143 | 107 | 111 | 85 | 85 | 203 | 203 | 114 | 116 | 125 | 125 |
| | GKB3 | 9 | 107 | 143 | 107 | 111 | 85 | 85 | 203 | 203 | 114 | 118 | 125 | 125 |
| | GKC2 | 9 | 107 | 143 | 107 | 111 | 85 | 85 | 203 | 203 | 114 | 118 | 125 | 125 |
| | GKC3 | 9 | 107 | 143 | 107 | 111 | 85 | 85 | 203 | 203 | 114 | 118 | 125 | 125 |
| | GKC4 | 9 | 107 | 143 | 107 | 111 | 85 | 85 | 203 | 203 | 114 | 118 | 125 | 125 |
| | GKD1 | 9 | 107 | 143 | 107 | 111 | 85 | 85 | 203 | 203 | 114 | 118 | 125 | 125 |
| | GKD2 | 9 | 107 | 143 | 107 | 111 | 85 | 85 | 203 | 203 | 114 | 118 | 125 | 125 |
| | GKD3 | 9 | 107 | 143 | 107 | 111 | 85 | 85 | 203 | 203 | 114 | 118 | 125 | 125 |
| | GKD4 | 9 | 107 | 143 | 107 | 111 | 85 | 85 | 203 | 203 | 100 | 118 | 125 | 127 |
| **Seedorf** | SEA1 | 11 | 89 | 116 | 107 | 109 | 87 | 87 | 200 | 227 | 114 | 120 | 127 | 129 |
| 3x3 m grid | SEA2 | 11 | 89 | 116 | 107 | 109 | 87 | 87 | 206 | 227 | 114 | 120 | 127 | 129 |
| | SEA3 | 11 | 89 | 116 | 107 | 109 | 87 | 87 | 200 | 227 | 114 | 120 | 127 | 129 |

| | | | | | | | | | | | | | | |
|---|---|---|---|---|---|---|---|---|---|---|---|---|---|---|
| | SEB1 | 11 | 89 | 116 | 107 | 109 | 87 | 87 | 200 | 227 | 114 | 120 | 127 | 129 |
| | SEB2 | 11 | 89 | 116 | 107 | 109 | 87 | 87 | 206 | 227 | 114 | 120 | 127 | 129 |
| | SEB3 | 11 | 89 | 116 | 107 | 109 | 87 | 87 | 200 | 227 | 114 | 120 | 127 | 129 |
| | SEB4 | 11 | 89 | 116 | 107 | 109 | 87 | 87 | 200 | 227 | 114 | 120 | 127 | 129 |
| | SEC1 | 11 | 89 | 116 | 107 | 109 | 87 | 87 | 206 | 227 | 114 | 122 | 127 | 129 |
| | SEC2 | 11 | 89 | 116 | 107 | 109 | 87 | 87 | 200 | 227 | 114 | 120 | 127 | 129 |
| | SEC3 | 11 | 89 | 116 | 107 | 109 | 87 | 87 | 200 | 227 | 114 | 120 | 127 | 129 |
| | SEC4 | 11 | 89 | 116 | 107 | 109 | 87 | 87 | 200 | 227 | 114 | 120 | 127 | 129 |
| | SED1 | 11 | 89 | 116 | 107 | 109 | 87 | 87 | 200 | 227 | 114 | 122 | 127 | 129 |
| | SED2 | 11 | 89 | 116 | 107 | 109 | 87 | 87 | 200 | 227 | 114 | 120 | 127 | 129 |
| | SED3 | 11 | 89 | 116 | 107 | 109 | 87 | 87 | 200 | 227 | 114 | 120 | 127 | 129 |
| | SED4 | 11 | 89 | 116 | 107 | 109 | 87 | 87 | 200 | 227 | 114 | 120 | 127 | 129 |
| **Restorf** | REA1 | 14 | 89 | 89 | 105 | 109 | 91 | 91 | 203 | 203 | 112 | 122 | 125 | 127 |
| 3x3 m grid | REA2 | 14 | 89 | 110 | 107 | 109 | 85 | 87 | 203 | 203 | 112 | 116 | 125 | 125 |
| | REA3 | 14 | 89 | 110 | 107 | 109 | 85 | 87 | 203 | 203 | 112 | 116 | 125 | 125 |
| | REA4 | 14 | 89 | 110 | 107 | 109 | 85 | 87 | 203 | 203 | 112 | 116 | 125 | 125 |
| | REB1 | 14 | 89 | 89 | 105 | 109 | 91 | 91 | 203 | 203 | 112 | 122 | 125 | 127 |
| | REB2 | 14 | 89 | 110 | 107 | 109 | 85 | 87 | 203 | 203 | 112 | 116 | 125 | 125 |
| | REB3 | 14 | 89 | 110 | 107 | 109 | 85 | 87 | 203 | 203 | 112 | 116 | 125 | 125 |
| | REB4 | 14 | 89 | 110 | 107 | 109 | 85 | 87 | 203 | 203 | 110 | 116 | 125 | 125 |
| | REC1 | 14 | 89 | 110 | 107 | 109 | 85 | 87 | 203 | 203 | 112 | 116 | 125 | 125 |
| | REC2 | 14 | 89 | 110 | 107 | 109 | 85 | 87 | 203 | 203 | 112 | 116 | 125 | 125 |
| | REC3 | 14 | 89 | 110 | 107 | 109 | 85 | 87 | 203 | 203 | 112 | 116 | 125 | 125 |
| | REC4 | 14 | 107 | 107 | 107 | 111 | 85 | 87 | 227 | 227 | 114 | 120 | 127 | 145 |
| | RED1 | 14 | 89 | 110 | 107 | 109 | 85 | 87 | 203 | 203 | 112 | 116 | 125 | 125 |
| | RED2 | 14 | 89 | 110 | 107 | 109 | 85 | 87 | 203 | 203 | 112 | 116 | 125 | 125 |
| | RED3 | 14 | 89 | 110 | 107 | 109 | 85 | 87 | 203 | 203 | 112 | 116 | 125 | 125 |
| | RED4 | 14 | 89 | 89 | 107 | 107 | 87 | 87 | 203 | 203 | 110 | 110 | 125 | 125 |
| **Restorf** | NRE-A1 | 141 | 89 | 89 | 105 | 109 | 91 | 91 | 203 | 203 | 112 | 120 | 125 | 127 |
| 10x10 m grid | NRE-A2 | 141 | 89 | 89 | 105 | 109 | 91 | 91 | 203 | 203 | 112 | 122 | 125 | 127 |
| | NRE-A6 | 141 | 104 | 113 | 105 | 109 | 85 | 87 | 203 | 203 | 108 | 112 | 125 | 125 |

**NSG Tonabgrabungen**
10x10 m grid

| | | | | | | | | | | | | | |
|---|---|---|---|---|---|---|---|---|---|---|---|---|---|
| NRE-B2 | 141 | 89 | 89 | 105 | 109 | 91 | 91 | 203 | 203 | 112 | 122 | 125 | 127 |
| NRE-B4 | 141 | 104 | 113 | 109 | 111 | 85 | 87 | 203 | 230 | 106 | 110 | 125 | 127 |
| NRE-B5 | 141 | 104 | 113 | 109 | 111 | 85 | 87 | 203 | 230 | 106 | 110 | 125 | 127 |
| NRE-C3 | 141 | 89 | 89 | 105 | 109 | 91 | 91 | 203 | 203 | 106 | 110 | 125 | 127 |
| NRE-C4 | 141 | 104 | 113 | 109 | 111 | 85 | 87 | 203 | 230 | 106 | 110 | 125 | 127 |
| NRE-C5 | 141 | 89 | 104 | 107 | 111 | 87 | 95 | 203 | 203 | 110 | 122 | 125 | 125 |
| NRE-C6 | 141 | 89 | 104 | 107 | 111 | 87 | 95 | 203 | 203 | 110 | 122 | 125 | 125 |
| NRE-D1 | 141 | 89 | 89 | 105 | 109 | 91 | 91 | 203 | 203 | 112 | 122 | 125 | 127 |
| NRE-D3 | 141 | 89 | 89 | 105 | 109 | 91 | 91 | 203 | 203 | 112 | 122 | 125 | 127 |
| NRE-D4 | 141 | 89 | 89 | 105 | 109 | 91 | 91 | 203 | 203 | 112 | 122 | 125 | 127 |
| NRE-D5 | 141 | 89 | 104 | 107 | 111 | 87 | 95 | 203 | 203 | 110 | 122 | 125 | 125 |
| NRE-E2 | 141 | 89 | 113 | 105 | 109 | 91 | 91 | 203 | 203 | 112 | 122 | 125 | 127 |
| NRE-E5 | 141 | 89 | 89 | 105 | 109 | 91 | 91 | 203 | 203 | 112 | 122 | 125 | 127 |
| NRE-F1 | 141 | 89 | 89 | 105 | 109 | 91 | 91 | 203 | 203 | 112 | 122 | 125 | 127 |
| NRE-F4 | 141 | 89 | 89 | 105 | 109 | 91 | 91 | 203 | 203 | 112 | 122 | 125 | 127 |
| NRE-F5 | 141 | 89 | 89 | 107 | 107 | 85 | 87 | 203 | 203 | 112 | 114 | 125 | 125 |
| NRE-F6 | 141 | 89 | 89 | 107 | 107 | 85 | 87 | 203 | 203 | 112 | 114 | 125 | 125 |
| NTO-A5 | 202 | 89 | 110 | 107 | 109 | 85 | 85 | 203 | 230 | 122 | 128 | 127 | 127 |
| NTO-A6 | 202 | 89 | 110 | 107 | 109 | 85 | 85 | 203 | 230 | 122 | 128 | 127 | 127 |
| NTO-B1 | 202 | 89 | 110 | 105 | 107 | 85 | 85 | 203 | 203 | 116 | 120 | 125 | 125 |
| NTO-B2 | 202 | 89 | 110 | 107 | 109 | 85 | 85 | 203 | 230 | 124 | 126 | 127 | 127 |
| NTO-B5 | 202 | 89 | 110 | 107 | 109 | 85 | 85 | 203 | 230 | 122 | 128 | 127 | 127 |
| NTO-B6 | 202 | 89 | 110 | 107 | 109 | 85 | 85 | 203 | 230 | 122 | 128 | 127 | 127 |
| NTO-C3 | 202 | 89 | 110 | 107 | 109 | 85 | 85 | 203 | 230 | 122 | 128 | 127 | 127 |
| NTO-C4 | 202 | 89 | 110 | 107 | 109 | 85 | 85 | 203 | 230 | 122 | 128 | 127 | 127 |
| NTO-C6 | 202 | 89 | 110 | 107 | 109 | 85 | 87 | 203 | 230 | 122 | 128 | 127 | 127 |
| NTO-D2 | 202 | 89 | 110 | 107 | 109 | 85 | 85 | 203 | 230 | 124 | 128 | 127 | 127 |
| NTO-D3 | 202 | 89 | 110 | 107 | 109 | 85 | 85 | 203 | 230 | 122 | 128 | 127 | 127 |
| NTO-D4 | 202 | 89 | 110 | 107 | 109 | 85 | 85 | 203 | 230 | 122 | 128 | 127 | 127 |
| NTO-E1 | 202 | 89 | 110 | 107 | 109 | 85 | 85 | 203 | 230 | 122 | 128 | 127 | 127 |
| NTO-E2 | 202 | 89 | 110 | 107 | 109 | 85 | 85 | 203 | 230 | 122 | 128 | 127 | 127 |
| NTO-E4 | 202 | 89 | 110 | 107 | 109 | 85 | 85 | 203 | 230 | 122 | 128 | 127 | 127 |

| | | | | | | | | | | | | | | |
|---|---|---|---|---|---|---|---|---|---|---|---|---|---|---|
| | NTO-E5 | 202 | 89 | 110 | 107 | 109 | 85 | 87 | 203 | 230 | 122 | 128 | 127 | 127 |
| | NTO-E6 | 202 | 89 | 110 | 107 | 109 | 85 | 85 | 203 | 230 | 122 | 128 | 127 | 127 |
| | NTO-F1 | 202 | 89 | 107 | 107 | 109 | 85 | 85 | 203 | 230 | 122 | 128 | 127 | 127 |
| | NTO-F4 | 202 | 89 | 110 | 107 | 109 | 85 | 85 | 203 | 224 | 122 | 128 | 127 | 127 |
| | NTO-F6 | 202 | 89 | 110 | 107 | 109 | 85 | 85 | 203 | 230 | 122 | 128 | 127 | 127 |
| **Osterburg** | OS-A1 | 24 | 110 | 125 | 107 | 111 | 89 | 91 | 224 | 227 | 110 | 110 | 125 | 125 |
| 3x3 m grid | OS-A2 | 24 | 110 | 125 | 107 | 111 | 89 | 91 | 224 | 227 | 110 | 110 | 125 | 125 |
| | OS-A3 | 24 | 110 | 125 | 107 | 111 | 89 | 91 | 224 | 227 | 110 | 110 | 125 | 125 |
| | OS-A4 | 24 | 110 | 125 | 107 | 111 | 89 | 91 | 224 | 227 | 110 | 110 | 125 | 125 |
| | OS-B2 | 24 | 110 | 125 | 107 | 111 | 89 | 91 | 224 | 227 | 110 | 110 | 125 | 125 |
| | OS-B3 | 24 | 110 | 125 | 107 | 111 | 89 | 91 | 224 | 227 | 110 | 110 | 125 | 125 |
| | OS-B4 | 24 | 110 | 125 | 107 | 111 | 89 | 91 | 224 | 227 | 110 | 110 | 125 | 125 |
| | OS-C1 | 24 | 110 | 125 | 107 | 111 | 89 | 91 | 224 | 227 | 110 | 110 | 125 | 125 |
| | OS-C3 | 24 | 110 | 125 | 107 | 111 | 89 | 91 | 224 | 227 | 110 | 110 | 125 | 125 |
| | OS-C4 | 24 | 110 | 125 | 107 | 111 | 89 | 91 | 224 | 227 | 110 | 110 | 125 | 127 |
| | OS-D3 | 24 | 110 | 125 | 107 | 111 | 89 | 91 | 224 | 227 | 110 | 110 | 125 | 127 |
| **Beuster Altarm** | BAA1 | 29 | 113 | 116 | 107 | 107 | 89 | 89 | 203 | 233 | 102 | 108 | 125 | 131 |
| 3x3 m grid | BAA2 | 29 | 113 | 116 | 107 | 107 | 89 | 89 | 203 | 233 | 102 | 108 | 125 | 131 |
| | BAA3 | 29 | 113 | 119 | 107 | 107 | 89 | 89 | 203 | 233 | 102 | 108 | 125 | 131 |
| | BAA4 | 29 | 113 | 119 | 107 | 107 | 89 | 89 | 203 | 233 | 102 | 108 | 125 | 131 |
| | BAB1 | 29 | 113 | 119 | 107 | 107 | 89 | 89 | 203 | 233 | 102 | 108 | 125 | 131 |
| | BAB2 | 29 | 113 | 119 | 107 | 107 | 89 | 89 | 203 | 233 | 102 | 108 | 125 | 131 |
| | BAB3 | 29 | 113 | 119 | 107 | 107 | 89 | 89 | 203 | 233 | 102 | 108 | 125 | 131 |
| | BAB4 | 29 | 113 | 119 | 107 | 107 | 89 | 89 | 203 | 233 | 102 | 108 | 125 | 131 |
| | BAC1 | 29 | 113 | 119 | 107 | 107 | 89 | 89 | 203 | 233 | 102 | 108 | 125 | 131 |
| | BAC2 | 29 | 113 | 119 | 107 | 107 | 89 | 89 | 203 | 233 | 102 | 108 | 125 | 131 |
| | BAC3 | 29 | 113 | 119 | 107 | 107 | 89 | 89 | 203 | 233 | 102 | 108 | 125 | 131 |
| | BAC4 | 29 | 113 | 119 | 107 | 107 | 89 | 89 | 203 | 233 | 102 | 108 | 125 | 131 |
| | BAD1 | 29 | 113 | 119 | 107 | 107 | 89 | 89 | 203 | 233 | 102 | 108 | 125 | 131 |
| | BAD2 | 29 | 113 | 119 | 107 | 107 | 89 | 89 | 203 | 233 | 102 | 108 | 125 | 131 |
| | BAD3 | 29 | 113 | 119 | 107 | 107 | 89 | 89 | 203 | 233 | 102 | 108 | 125 | 131 |

| | | | | | | | | | | | | | | |
|---|---|---|---|---|---|---|---|---|---|---|---|---|---|---|
| | BAD4 | 29 | 113 | 119 | 107 | 107 | 89 | 89 | 203 | 233 | 102 | 108 | 125 | 131 |
| **Fischbeck Deich** 3x3 m grid | FDA1 | 31 | 122 | 134 | 107 | 109 | 85 | 87 | 206 | 230 | 100 | 126 | 127 | 127 |
| | FDA2 | 31 | 122 | 134 | 107 | 109 | 85 | 87 | 206 | 230 | 100 | 126 | 127 | 127 |
| | FDA3 | 31 | 122 | 134 | 107 | 109 | 85 | 87 | 206 | 230 | 100 | 126 | 127 | 127 |
| | FDA4 | 31 | 122 | 134 | 107 | 109 | 85 | 87 | 206 | 230 | 100 | 126 | 127 | 127 |
| | FDB1 | 31 | 122 | 134 | 107 | 109 | 85 | 87 | 206 | 230 | 100 | 126 | 127 | 127 |
| | FDB2 | 31 | 122 | 134 | 107 | 109 | 85 | 87 | 206 | 230 | 100 | 126 | 127 | 127 |
| | FDB3 | 31 | 122 | 134 | 107 | 109 | 85 | 87 | 206 | 230 | 100 | 126 | 127 | 127 |
| | FDB4 | 31 | 122 | 134 | 107 | 109 | 85 | 87 | 206 | 230 | 100 | 126 | 127 | 127 |
| | FDC1 | 31 | 122 | 134 | 107 | 109 | 85 | 87 | 206 | 230 | 100 | 126 | 127 | 127 |
| | FDC2 | 31 | 122 | 134 | 107 | 109 | 85 | 87 | 206 | 230 | 100 | 126 | 127 | 127 |
| | FDC3 | 31 | 122 | 134 | 107 | 109 | 85 | 87 | 206 | 233 | 100 | 126 | 127 | 127 |
| | FDC4 | 31 | 122 | 134 | 107 | 109 | 85 | 87 | 206 | 230 | 100 | 126 | 127 | 127 |
| | FDD1 | 31 | 122 | 134 | 107 | 109 | 85 | 87 | 206 | 230 | 100 | 126 | 127 | 127 |
| | FDD2 | 31 | 122 | 134 | 107 | 109 | 85 | 87 | 206 | 230 | 100 | 126 | 127 | 127 |
| | FDD3 | 31 | 122 | 134 | 107 | 109 | 85 | 87 | 206 | 230 | 100 | 126 | 127 | 127 |
| | FDD4 | 31 | 122 | 134 | 107 | 109 | 85 | 87 | 206 | 230 | 100 | 126 | 127 | 127 |
| **Bucher Brack** 3x3 m grid | BBA1 | 32 | 98 | 110 | 105 | 111 | 87 | 91 | 194 | 206 | 112 | 112 | 127 | 129 |
| | BBA3 | 32 | 116 | 137 | 109 | 111 | 85 | 87 | 203 | 203 | 102 | 120 | 125 | 127 |
| | BBA4 | 32 | 98 | 110 | 105 | 111 | 87 | 91 | 194 | 206 | 112 | 112 | 127 | 129 |
| | BBB1 | 32 | 98 | 110 | 105 | 111 | 87 | 91 | 194 | 206 | 112 | 112 | 127 | 129 |
| | BBB2 | 32 | 98 | 110 | 105 | 111 | 87 | 91 | 194 | 206 | 112 | 112 | 127 | 129 |
| | BBB3 | 32 | 98 | 110 | 105 | 111 | 87 | 91 | 194 | 206 | 112 | 112 | 127 | 129 |
| | BBB4 | 32 | 113 | 137 | 109 | 111 | 85 | 87 | 203 | 203 | 102 | 120 | 125 | 127 |
| | BBC1 | 32 | 98 | 110 | 105 | 111 | 87 | 91 | 194 | 206 | 112 | 112 | 127 | 129 |
| | BBC2 | 32 | 98 | 110 | 105 | 111 | 87 | 91 | 194 | 206 | 112 | 112 | 127 | 129 |
| | BBC3 | 32 | 98 | 110 | 105 | 111 | 87 | 91 | 194 | 206 | 112 | 112 | 127 | 129 |
| | BBD1 | 32 | 98 | 110 | 105 | 111 | 87 | 91 | 194 | 206 | 112 | 112 | 127 | 129 |
| | BBD2 | 32 | 98 | 110 | 105 | 111 | 87 | 91 | 194 | 206 | 112 | 112 | 127 | 129 |
| | BBD3 | 32 | 98 | 107 | 107 | 107 | 85 | 91 | 194 | 227 | 112 | 122 | 125 | 127 |
| | BBD4 | 32 | 113 | 137 | 109 | 111 | 85 | 87 | 203 | 203 | 102 | 120 | 125 | 127 |

**Wulfener Bruch**
3x3 m grid

| | | | | | | | | | | | | | |
|---|---|---|---|---|---|---|---|---|---|---|---|---|---|
| WB-A1 | 38 | 89 | 89 | 105 | 107 | 85 | 87 | 203 | 206 | 102 | 120 | 125 | 125 |
| WB-A2 | 38 | 89 | 89 | 105 | 107 | 85 | 87 | 203 | 206 | 102 | 122 | 125 | 125 |
| WB-A3 | 38 | 89 | 89 | 105 | 107 | 85 | 87 | 203 | 206 | 102 | 122 | 125 | 125 |
| WB-A4 | 38 | 89 | 89 | 105 | 107 | 85 | 87 | 203 | 206 | 102 | 120 | 125 | 125 |
| WB-B1 | 38 | 89 | 89 | 105 | 107 | 85 | 87 | 203 | 206 | 102 | 122 | 125 | 125 |
| WB-B2 | 38 | 89 | 89 | 105 | 107 | 85 | 87 | 203 | 206 | 102 | 122 | 125 | 125 |
| WB-B3 | 38 | 89 | 89 | 105 | 107 | 85 | 87 | 203 | 206 | 102 | 122 | 125 | 125 |
| WB-B4 | 38 | 89 | 89 | 105 | 107 | 85 | 87 | 203 | 206 | 102 | 122 | 125 | 127 |
| WB-C1 | 38 | 89 | 89 | 105 | 107 | 85 | 87 | 203 | 206 | 102 | 122 | 125 | 125 |
| WB-C2 | 38 | 89 | 89 | 105 | 107 | 85 | 87 | 203 | 206 | 102 | 122 | 125 | 125 |
| WB-C3 | 38 | 89 | 89 | 105 | 107 | 85 | 87 | 203 | 206 | 102 | 122 | 125 | 125 |
| WB-C4 | 38 | 89 | 89 | 105 | 107 | 85 | 87 | 203 | 206 | 102 | 122 | 125 | 125 |
| WB-D1 | 38 | 89 | 89 | 105 | 107 | 85 | 87 | 203 | 206 | 102 | 122 | 125 | 125 |
| WB-D2 | 38 | 89 | 89 | 105 | 107 | 85 | 87 | 203 | 206 | 102 | 122 | 125 | 125 |
| WB-D3 | 38 | 89 | 89 | 105 | 107 | 85 | 87 | 203 | 206 | 102 | 122 | 125 | 125 |
| WB-D4 | 38 | 89 | 89 | 105 | 107 | 85 | 87 | 203 | 206 | 102 | 122 | 125 | 125 |

**NE of Aken**
3x3 m grid

| | | | | | | | | | | | | | |
|---|---|---|---|---|---|---|---|---|---|---|---|---|---|
| AK-A1 | 39 | 89 | 89 | 107 | 109 | 85 | 87 | 203 | 212 | 100 | 122 | 125 | 127 |
| AK-A2 | 39 | 89 | 89 | 107 | 109 | 85 | 87 | 203 | 212 | 122 | 122 | 125 | 127 |
| AK-A3 | 39 | 89 | 89 | 107 | 109 | 85 | 87 | 203 | 212 | 122 | 122 | 125 | 127 |
| AK-A4 | 39 | 89 | 89 | 107 | 109 | 85 | 87 | 203 | 212 | 122 | 122 | 125 | 127 |
| AK-B1 | 39 | 89 | 89 | 107 | 109 | 85 | 87 | 203 | 212 | 122 | 122 | 125 | 127 |
| AK-B2 | 39 | 89 | 89 | 107 | 109 | 85 | 87 | 203 | 212 | 122 | 122 | 125 | 127 |
| AK-B3 | 39 | 89 | 89 | 107 | 109 | 85 | 87 | 203 | 212 | 122 | 122 | 125 | 127 |
| AK-B4 | 39 | 89 | 89 | 107 | 109 | 87 | 87 | 203 | 212 | 122 | 122 | 125 | 127 |
| AK-C1 | 39 | 89 | 89 | 107 | 109 | 85 | 87 | 203 | 212 | 122 | 122 | 125 | 127 |
| AK-C2 | 39 | 89 | 89 | 107 | 109 | 85 | 87 | 203 | 212 | 122 | 122 | 125 | 127 |
| AK-C3 | 39 | 89 | 89 | 107 | 109 | 87 | 87 | 203 | 212 | 122 | 122 | 125 | 127 |
| AK-C4 | 39 | 89 | 89 | 107 | 109 | 85 | 87 | 203 | 212 | 122 | 122 | 125 | 127 |
| AK-D1 | 39 | 89 | 89 | 107 | 109 | 85 | 87 | 203 | 212 | 122 | 122 | 125 | 127 |
| AK-D2 | 39 | 89 | 89 | 107 | 109 | 85 | 87 | 203 | 212 | 122 | 122 | 125 | 127 |
| AK-D3 | 39 | 89 | 89 | 107 | 109 | 85 | 87 | 203 | 212 | 122 | 122 | 125 | 127 |

**Wiesen am Schwedenhaus**
3x3 m grid

| | | | | | | | | | | | | | |
|---|---|---|---|---|---|---|---|---|---|---|---|---|---|
| SW-A1 | 40 | 89 | 122 | 105 | 111 | 89 | 91 | 206 | 209 | 110 | 112 | 127 | 127 |
| SW-A2 | 40 | 89 | 122 | 105 | 111 | 89 | 91 | 206 | 209 | 110 | 112 | 127 | 127 |
| SW-A3 | 40 | 89 | 122 | 105 | 111 | 89 | 91 | 206 | 209 | 110 | 112 | 127 | 127 |
| SW-A4 | 40 | 89 | 122 | 105 | 111 | 89 | 91 | 206 | 209 | 110 | 112 | 127 | 127 |
| SW-B1 | 40 | 89 | 122 | 105 | 111 | 89 | 91 | 206 | 209 | 110 | 112 | 127 | 127 |
| SW-B2 | 40 | 89 | 122 | 105 | 111 | 89 | 91 | 206 | 209 | 110 | 112 | 127 | 127 |
| SW-B3 | 40 | 89 | 122 | 105 | 111 | 89 | 91 | 206 | 209 | 110 | 112 | 127 | 127 |
| SW-B4 | 40 | 89 | 122 | 105 | 111 | 89 | 91 | 206 | 209 | 110 | 112 | 127 | 129 |
| SW-C1 | 40 | 89 | 122 | 105 | 111 | 89 | 91 | 206 | 209 | 110 | 112 | 127 | 127 |
| SW-C2 | 40 | 89 | 122 | 105 | 111 | 89 | 91 | 206 | 209 | 110 | 112 | 127 | 127 |
| SW-C3 | 40 | 89 | 122 | 105 | 111 | 89 | 91 | 206 | 209 | 110 | 112 | 127 | 129 |
| SW-C4 | 40 | 89 | 122 | 105 | 111 | 89 | 91 | 206 | 209 | 110 | 112 | 127 | 129 |
| SW-D1 | 40 | 89 | 122 | 105 | 111 | 89 | 91 | 206 | 209 | 110 | 112 | 127 | 127 |
| SW-D2 | 40 | 89 | 122 | 105 | 111 | 89 | 91 | 206 | 209 | 110 | 112 | 127 | 127 |
| SW-D3 | 40 | 89 | 122 | 105 | 111 | 89 | 91 | 206 | 209 | 110 | 112 | 127 | 129 |
| SW-D4 | 40 | 89 | 122 | 105 | 111 | 89 | 91 | 206 | 209 | 110 | 112 | 127 | 129 |

**S of Rosslau**
3x3 m grid

| | | | | | | | | | | | | | |
|---|---|---|---|---|---|---|---|---|---|---|---|---|---|
| RL-A1 | 41 | 101 | 116 | 105 | 109 | 87 | 91 | 203 | 203 | 106 | 112 | 125 | 127 |
| RL-A2 | 41 | 101 | 116 | 105 | 109 | 87 | 91 | 203 | 203 | 106 | 112 | 125 | 127 |
| RL-A3 | 41 | 101 | 116 | 105 | 109 | 87 | 91 | 203 | 203 | 106 | 112 | 125 | 127 |
| RL-A4 | 41 | 101 | 116 | 105 | 109 | 87 | 91 | 203 | 203 | 106 | 112 | 125 | 127 |
| RL-B1 | 41 | 101 | 116 | 105 | 109 | 87 | 91 | 203 | 203 | 106 | 112 | 125 | 127 |
| RL-B2 | 41 | 101 | 116 | 105 | 109 | 87 | 91 | 203 | 203 | 106 | 112 | 125 | 127 |
| RL-B3 | 41 | 101 | 116 | 105 | 109 | 87 | 91 | 203 | 203 | 106 | 112 | 125 | 127 |
| RL-B4 | 41 | 101 | 116 | 105 | 109 | 87 | 91 | 203 | 203 | 106 | 112 | 125 | 127 |
| RL-C1 | 41 | 101 | 116 | 105 | 109 | 87 | 91 | 203 | 203 | 106 | 112 | 125 | 127 |
| RL-C2 | 41 | 101 | 116 | 105 | 109 | 87 | 91 | 203 | 203 | 106 | 112 | 125 | 127 |
| RL-C3 | 41 | 101 | 116 | 105 | 109 | 87 | 91 | 203 | 203 | 106 | 112 | 125 | 127 |
| RL-C4 | 41 | 101 | 116 | 105 | 109 | 87 | 91 | 203 | 203 | 106 | 112 | 125 | 127 |
| RL-D1 | 41 | 101 | 116 | 105 | 109 | 87 | 91 | 203 | 203 | 106 | 114 | 125 | 127 |
| RL-D2 | 41 | 101 | 116 | 105 | 109 | 87 | 91 | 203 | 203 | 106 | 112 | 125 | 127 |
| RL-D3 | 41 | 101 | 116 | 105 | 109 | 87 | 91 | 203 | 203 | 106 | 112 | 125 | 127 |

| | | | | | | | | | | | | | | |
|---|---|---|---|---|---|---|---|---|---|---|---|---|---|---|
| | RL-D4 | 41 | 101 | 116 | 105 | 109 | 87 | 91 | 203 | 203 | 106 | 112 | 125 | 127 |
| **E of Vockerode** | VO-A1 | 44 | 89 | 89 | 107 | 109 | 87 | 87 | 197 | 203 | 112 | 112 | 125 | 127 |
| 3x3 m grid | VO-A3 | 44 | 89 | 89 | 105 | 109 | 85 | 87 | 203 | 203 | 100 | 100 | 125 | 125 |
| | VO-A4 | 44 | 89 | 89 | 105 | 109 | 85 | 87 | 203 | 203 | 100 | 100 | 125 | 125 |
| | VO-B2 | 44 | 89 | 89 | 105 | 109 | 85 | 87 | 203 | 203 | 100 | 100 | 125 | 125 |
| | VO-B3 | 44 | 89 | 89 | 105 | 109 | 85 | 87 | 203 | 203 | 100 | 100 | 125 | 125 |
| | VO-B4 | 44 | 89 | 89 | 105 | 109 | 85 | 87 | 203 | 203 | 100 | 100 | 125 | 127 |
| | VO-C1 | 44 | 89 | 89 | 105 | 109 | 85 | 87 | 203 | 203 | 100 | 100 | 125 | 125 |
| | VO-C2 | 44 | 89 | 89 | 105 | 109 | 85 | 87 | 203 | 203 | 100 | 100 | 125 | 127 |
| | VO-C3 | 44 | 89 | 89 | 105 | 109 | 85 | 87 | 203 | 203 | 100 | 100 | 125 | 125 |
| | VO-C4 | 44 | 89 | 89 | 105 | 109 | 85 | 87 | 203 | 203 | 100 | 100 | 125 | 127 |
| | VO-D1 | 44 | 89 | 89 | 105 | 109 | 85 | 87 | 203 | 203 | 100 | 100 | 125 | 127 |
| | VO-D2 | 44 | 89 | 89 | 105 | 109 | 85 | 87 | 203 | 203 | 100 | 100 | 125 | 127 |
| | VO-D3 | 44 | 89 | 89 | 105 | 109 | 85 | 87 | 203 | 203 | 100 | 100 | 125 | 125 |
| | VO-D4 | 44 | 89 | 89 | 105 | 109 | 85 | 87 | 203 | 203 | 100 | 100 | 125 | 127 |
| **E of Vockerode** | NVO-A1 | 444 | 89 | 113 | 107 | 107 | 85 | 87 | 206 | 227 | 116 | 130 | 125 | 125 |
| 10x10 m grid | NVO-A3 | 444 | 89 | 113 | 107 | 107 | 85 | 87 | 206 | 227 | 116 | 130 | 125 | 125 |
| | NVO-A4 | 444 | 89 | 113 | 107 | 107 | 85 | 87 | 206 | 227 | 116 | 130 | 125 | 125 |
| | NVO-A5 | 444 | 89 | 113 | 107 | 107 | 85 | 87 | 206 | 227 | 116 | 132 | 125 | 125 |
| | NVO-B1 | 444 | 89 | 113 | 107 | 107 | 85 | 87 | 206 | 227 | 116 | 130 | 125 | 125 |
| | NVO-B2 | 444 | 89 | 113 | 107 | 107 | 85 | 87 | 206 | 227 | 116 | 130 | 125 | 125 |
| | NVO-B3 | 444 | 89 | 113 | 107 | 107 | 85 | 87 | 206 | 227 | 116 | 130 | 125 | 125 |
| | NVO-B4 | 444 | 89 | 113 | 107 | 107 | 85 | 87 | 206 | 227 | 116 | 130 | 125 | 125 |
| | NVO-B5 | 444 | 89 | 113 | 107 | 107 | 85 | 87 | 206 | 227 | 116 | 130 | 125 | 125 |
| | NVO-C1 | 444 | 98 | 113 | 105 | 107 | 87 | 89 | 203 | 203 | 106 | 116 | 125 | 127 |
| | NVO-C2 | 444 | 98 | 113 | 105 | 107 | 87 | 89 | 203 | 203 | 100 | 114 | 125 | 127 |
| | NVO-C3 | 444 | 98 | 113 | 105 | 107 | 87 | 89 | 203 | 203 | 100 | 114 | 125 | 127 |
| | NVO-C4 | 444 | 89 | 113 | 107 | 107 | 85 | 87 | 206 | 227 | 116 | 130 | 125 | 125 |
| | NVO-C5 | 444 | 89 | 113 | 107 | 107 | 85 | 87 | 206 | 227 | 116 | 130 | 125 | 125 |
| | NVO-D1 | 444 | 101 | 113 | 105 | 107 | 87 | 89 | 203 | 203 | 106 | 116 | 125 | 127 |
| | NVO-D2 | 444 | 98 | 113 | 105 | 107 | 87 | 89 | 203 | 203 | 100 | 114 | 125 | 127 |

| | | | | | | | | | | | | | | |
|---|---|---|---|---|---|---|---|---|---|---|---|---|---|---|
| | NVO-D3 | 444 | 98 | 113 | 105 | 107 | 87 | 89 | 203 | 203 | 100 | 114 | 125 | 127 |
| | NVO-D4 | 444 | 89 | 113 | 107 | 107 | 85 | 87 | 206 | 227 | 116 | 130 | 125 | 125 |
| | NVO-E1 | 444 | 98 | 113 | 105 | 107 | 87 | 89 | 203 | 203 | 100 | 114 | 125 | 127 |
| | NVO-E2 | 444 | 89 | 134 | 109 | 111 | 87 | 89 | 203 | 203 | 114 | 114 | 125 | 127 |
| | NVO-E4 | 444 | 98 | 113 | 105 | 107 | 87 | 89 | 203 | 203 | 100 | 114 | 125 | 127 |
| | NVO-F0 | 444 | 89 | 113 | 107 | 107 | 85 | 87 | 206 | 227 | 116 | 130 | 125 | 125 |
| | NVO-Z4 | 444 | 98 | 113 | 105 | 107 | 87 | 89 | 203 | 203 | 106 | 116 | 125 | 127 |
| **E of Elster** | EL-A1 | 47 | 113 | 140 | 105 | 111 | 85 | 85 | 206 | 221 | 110 | 110 | 127 | 129 |
| 3x3 m grid | EL-A2 | 47 | 113 | 140 | 105 | 111 | 85 | 85 | 206 | 221 | 110 | 110 | 127 | 129 |
| | EL-A3 | 47 | 113 | 140 | 105 | 111 | 85 | 85 | 206 | 221 | 110 | 110 | 127 | 129 |
| | EL-A4 | 47 | 113 | 206 | 105 | 105 | 85 | 85 | 203 | 203 | 112 | 124 | 125 | 133 |
| | EL-B1 | 47 | 113 | 206 | 105 | 105 | 85 | 85 | 203 | 203 | 112 | 124 | 125 | 133 |
| | EL-B2 | 47 | 113 | 206 | 105 | 105 | 85 | 85 | 203 | 203 | 112 | 124 | 125 | 133 |
| | EL-B3 | 47 | 113 | 206 | 105 | 105 | 85 | 85 | 203 | 203 | 112 | 124 | 125 | 133 |
| | EL-B4 | 47 | 113 | 203 | 105 | 105 | 85 | 85 | 203 | 203 | 112 | 124 | 125 | 133 |
| | EL-C1 | 47 | 113 | 206 | 105 | 105 | 85 | 85 | 203 | 203 | 112 | 124 | 125 | 133 |
| | EL-C2 | 47 | 113 | 206 | 105 | 105 | 85 | 85 | 203 | 203 | 112 | 124 | 125 | 133 |
| | EL-C3 | 47 | 113 | 206 | 105 | 105 | 85 | 85 | 203 | 203 | 112 | 124 | 125 | 133 |
| | EL-C4 | 47 | 113 | 206 | 105 | 105 | 85 | 85 | 203 | 203 | 112 | 124 | 127 | 133 |
| | EL-D1 | 47 | 113 | 206 | 105 | 105 | 85 | 85 | 203 | 203 | 112 | 124 | 125 | 133 |
| | EL-D2 | 47 | 113 | 206 | 105 | 105 | 85 | 85 | 203 | 203 | 112 | 124 | 125 | 133 |
| | EL-D3 | 47 | 113 | 203 | 105 | 105 | 85 | 85 | 203 | 203 | 112 | 124 | 125 | 133 |
| | EL-D4 | 47 | 113 | 206 | 105 | 105 | 85 | 85 | 203 | 203 | 112 | 124 | 125 | 133 |
