## Supplement S3 for "*Predominance of clonal propagation* conceals extinction risks of the highly endangered floodplain herb *Cnidium dubium*"

Supplement S3 – Results of the population genetic structure analysis

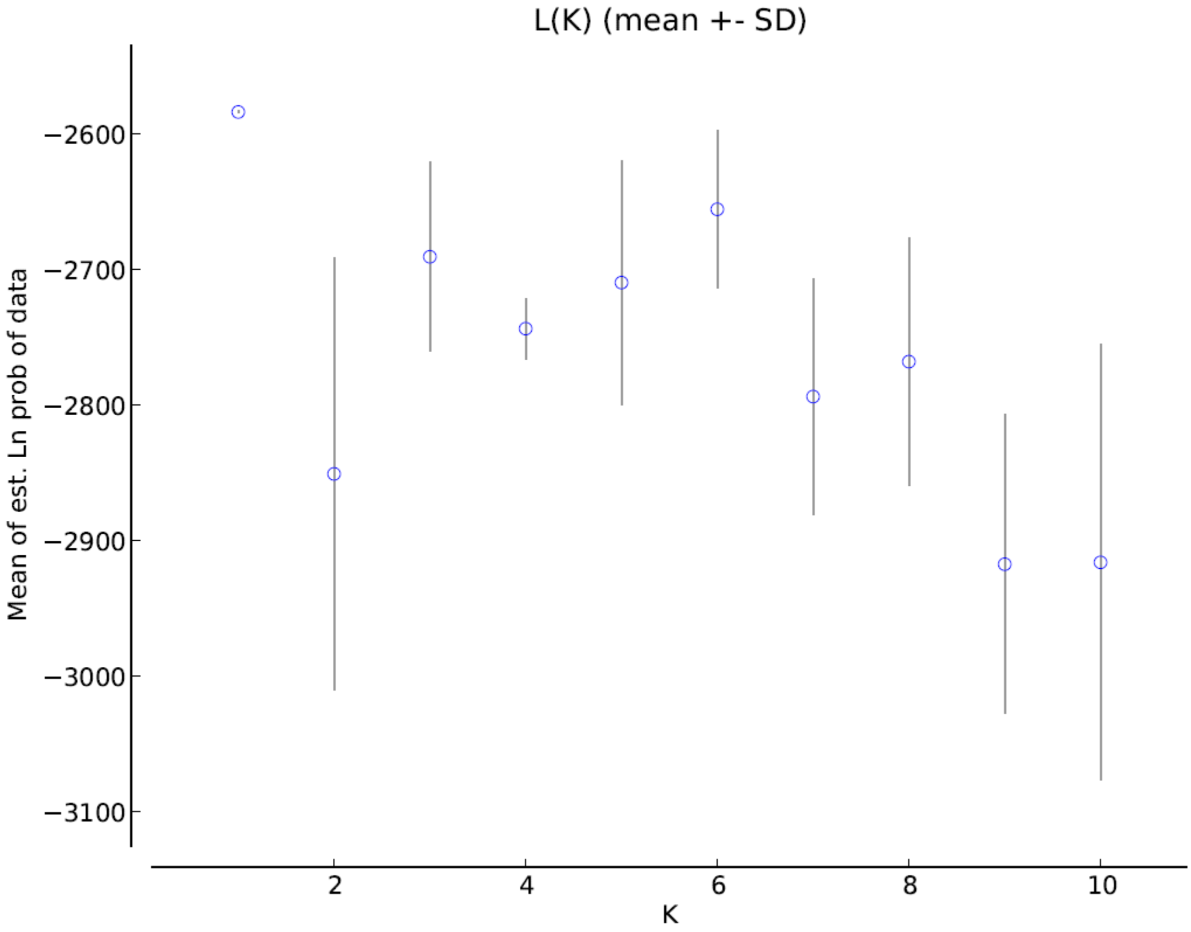


*Figure S1: Plot of mean estimated probability over K for 5 iterations per K.*

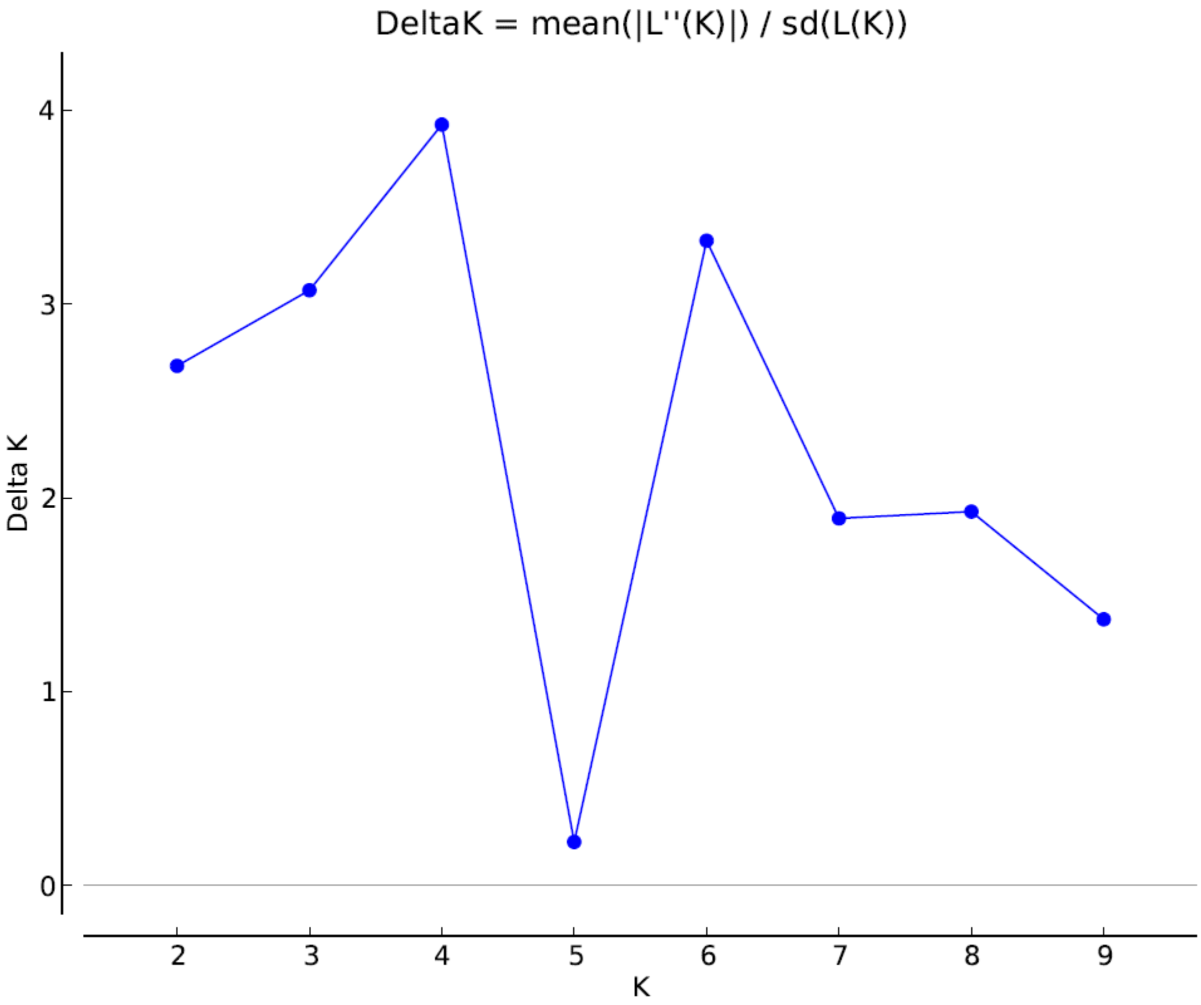


*Figure S2: Plot of deltaK over K.*

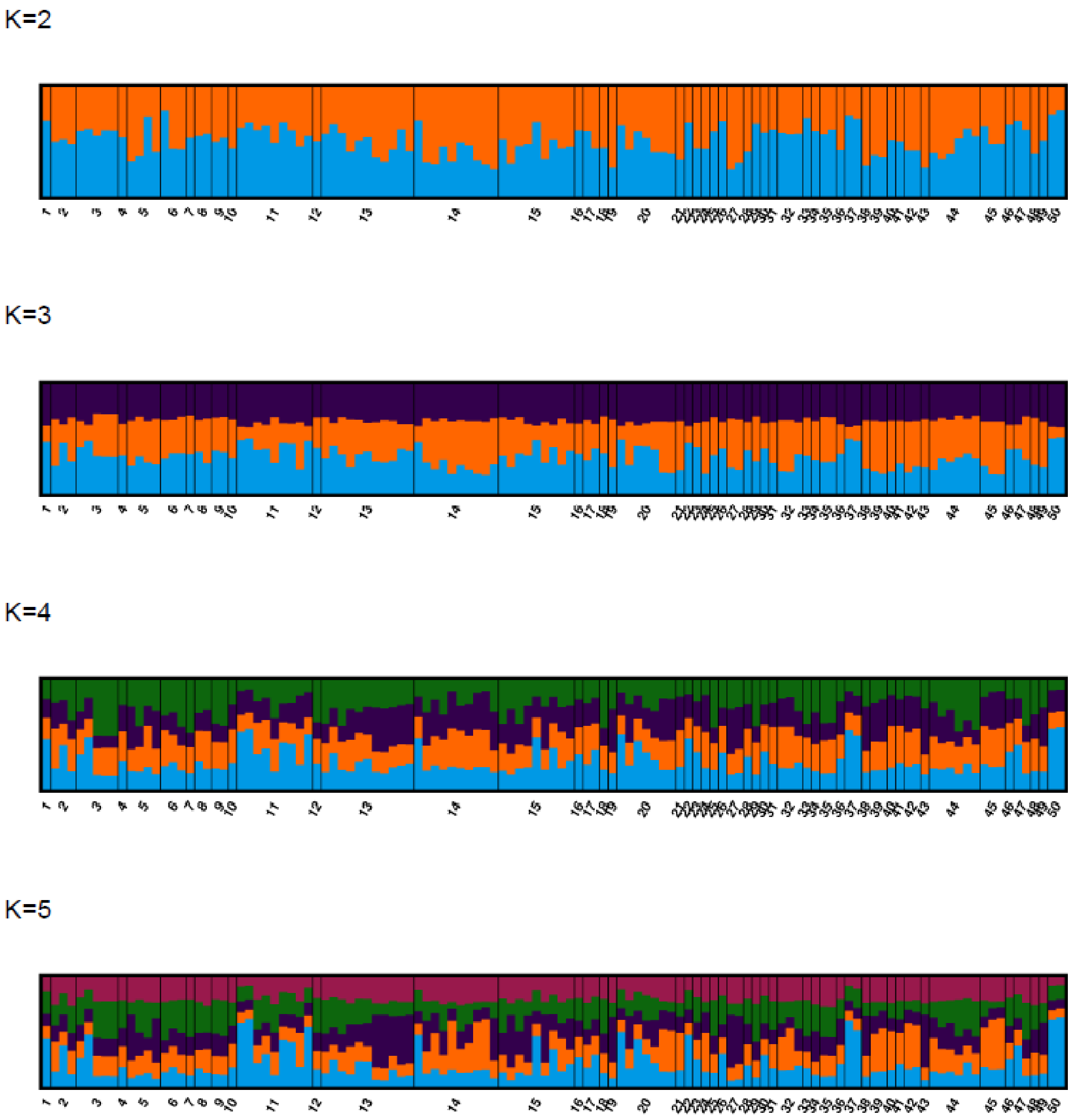


*Figure S3: STRUCTURE barplots for K = 2 -5 for 121 unique genotypes from 50 locations of C. dubium along the studied Elbe river stretch.*
